# Processed pseudogenes as dynamic substrates of vertebrate genome evolution

**DOI:** 10.64898/2026.06.26.734816

**Authors:** Rafael L V Mercuri, Gabriela D A Guardia, Daniela Moreira Mombach, Nathália Da Roz D’Alessandre, Helena Beatriz da Conceição, Thodoris Danis, Gabriel Arantes dos Santos, Filipe Ferreira dos Santos, Rafaela Carolina dos Anjos Schmidt, Matheus Pinheiro Moratto de Castro, Lorraine Christine de Oliveira, Alexander Birbrair, Marco Sollitto, Mary J. O’Connell, Antonis Rokas, Giulio Formenti, Marcela Uliano-Silva, Pedro A F Galante

**Author notes:** Contact author: Pedro A F Galante.

## Abstract

Pseudogenes, gene copies presumed nonfunctional, are widespread products of genome evolution, yet their retention under selection and functional significance across vertebrate diversity remain poorly understood. Here, by analyzing 244 high-quality, chromosome-scale genomes from the Vertebrate Genomes Project spanning seven major vertebrate lineages, we show that the most abundant class of pseudogenes, processed pseudogenes (retrocopies), is a dynamic substrate for evolutionary innovation rather than an inert relic. Retrocopy abundance varies by more than an order of magnitude across lineages, closely tracks autonomous retrotransposon content, and a considerable fraction retains intact open reading frames under purifying selection. We establish a two-stage model in which a conserved formation bias toward highly expressed housekeeping genes is followed by lineage-specific selective filtering that shapes distinct functional repertoires. Testing this model, we show that the mammalian X chromosome exports retrocopies to autosomes at significantly elevated rates enriched for functionally constrained copies, establishing meiotic sex chromosome inactivation as the selective driver. Furthermore, tumor suppressor gene retrocopies accumulate preferentially over oncogene retrocopies in large-bodied and long-lived mammalian lineages, identifying retrocopy-mediated tumor suppressor dosage expansion as a previously unrecognized genomic correlate of Peto’s paradox. Beyond cancer-related dynamics, retrocopy abundance itself correlates with key mammalian life-history traits, including brain mass, generation length, and reproductive timing, which suggests that retrocopy turnover is broadly coupled to organismal pace-of-life. These findings recast retrocopies as a major axis of vertebrate genome evolution and provide a comprehensive resource for studying gene duplicate innovation.

## MAIN

Gene duplication is a continuous, pervasive process in vertebrate genome evolution^1^, resulting in gene copies that span a spectrum from preserved paralogues to sequences classified as pseudogenes due to disrupted protein-coding potential^2,3^. Although traditionally regarded as nonfunctional relics of gene decay^1,3^, accumulating evidence shows that many loci annotated as pseudogenes are transcribed, contribute to the regulation of parental gene expression, or have been recruited into novel coding and non-coding roles^4–6^. The systematic extent of such functional contribution of pseudogenes across vertebrate diversity remains unresolved.

Pseudogenes arise through two mechanistically distinct routes^7^. Processed pseudogenes (retrocopies) form when spliced mRNAs are reverse-transcribed by retrotransposon machinery, predominantly Long Interspersed Nuclear Element-1 (LINE-1 or L1) in mammals, and inserted at new genomic locations as intronless copies that typically lack parental promoters^8,9^. Unprocessed pseudogenes (DNA-pseudogenes) derive instead from segmental, tandem, or whole-genome duplications and retain introns and ancestral (parental gene) regulatory elements^7^. Thus, because retrocopies enter new genomic environments stripped of inherited regulatory elements, any subsequent expression, retention of an open reading frame (ORF), or selective constraint must reflect *de novo* recruitment at the insertion site^3,4^, making them uniquely valuable substrates for studying gene innovation. The functional relevance of such recruitment is exemplified by *PTENP1*, a *PTEN* retrocopy that modulates parental gene abundance through shared microRNAs and influences oncogenic phenotypes^4^, and by *GLUD2*, a primate-specific retrocopy with neural-tissue function and adaptive substitutions of their parental gene (*GLUD*) function in glutamate metabolism^10^.

Despite these established cases, systematic characterization of retrocopies has been limited by the scarcity of chromosome-scale, standardized genome assemblies across vertebrate phylogeny^11^. Completion of Phase I of the Vertebrate Genomes Project (VGP) overcomes this limitation, providing high-quality assemblies and uniform annotations for hundreds of species^12^.

Here, we characterize retrocopies and DNA-pseudogenes across 244 species spanning seven major vertebrate lineages. Retrocopies outnumber DNA-pseudogenes by more than threefold and constitute the predominant pseudogene class. Their abundance varies by more than an order of magnitude across lineages, tracks autonomous non-long terminal repeat (non-LTR) retrotransposon content, and a substantial fraction retains intact ORFs under purifying selection. We establish a two-stage model of retrocopy evolution: a conserved formation bias driven by germline transcriptional availability, followed by lineage-specific selective filtering of the retained retrocopy repertoire. We test this model across three domains. First, the mammalian X chromosome exports retrocopies to autosomes at elevated rates and with enrichment for conserved and coding retrocopies under purifying selection, a pattern absent from the avian ZW system. Second, cancer-associated genes are pervasive retroduplication substrates, with tumor suppressor retrocopies accumulating preferentially over oncogene retrocopies in large-bodied and long-lived mammalian lineages, identifying retrocopy-mediated tumor suppressor dosage expansion as a previously unrecognized genomic correlate of Peto’s paradox. Third, retrocopy variation correlates with mammalian life-history traits, including brain mass, generation length, and reproductive timing. These findings establish retrocopies as a major axis of vertebrate genome evolution and provide a comprehensive resource for the study of gene duplicate innovation.

## RESULTS

### Retrocopies dominate vertebrate pseudogene landscapes and track non-LTR retrotransposon content

Across 244 chromosome-scale VGP genomes with gene annotations (2025 Phase I data freeze^12^ spanning seven major vertebrate lineages (**Fig. 1a**; **Supplementary Table 1**), we identified 555,947 retrocopies using RCPedia^13^ and 179,477 DNA-pseudogenes from curated RefSeq annotations^14^, corresponding to 9.7% and 3.1% of all annotated loci, respectively (**Fig. 1b-c**; **Supplementary Table 2**). Retrocopies were the predominant pseudogene class in five of seven lineages (**Extended Data Fig. 1a**), exceeding DNA-pseudogenes by more than threefold genome-wide (**Fig 1c**). Mammals carried the highest fraction (21.6%; 478,469 retrocopies), followed by amphibians (6.1%; 21,861) and cartilaginous fishes (5.4%; 14,081), with birds at the opposite extreme (0.8%; 7,263) and ray-finned fishes (1.4%; 22,128) showing a DNA-pseudogene-biased profile consistent with the legacy of teleost-specific whole-genome duplication^15^. Both the retrocopy-to-parental ratio (∼2.8 in mammals to ∼1.5 in lepidosaurs; **Supplementary Fig. 1a**) and the median retrocopy length (375 bp in amphibians to 674 bp in mammals; **Supplementary Fig. 1b**) varied across lineages, underscoring divergent retroduplication dynamics.

**Fig. 1.**
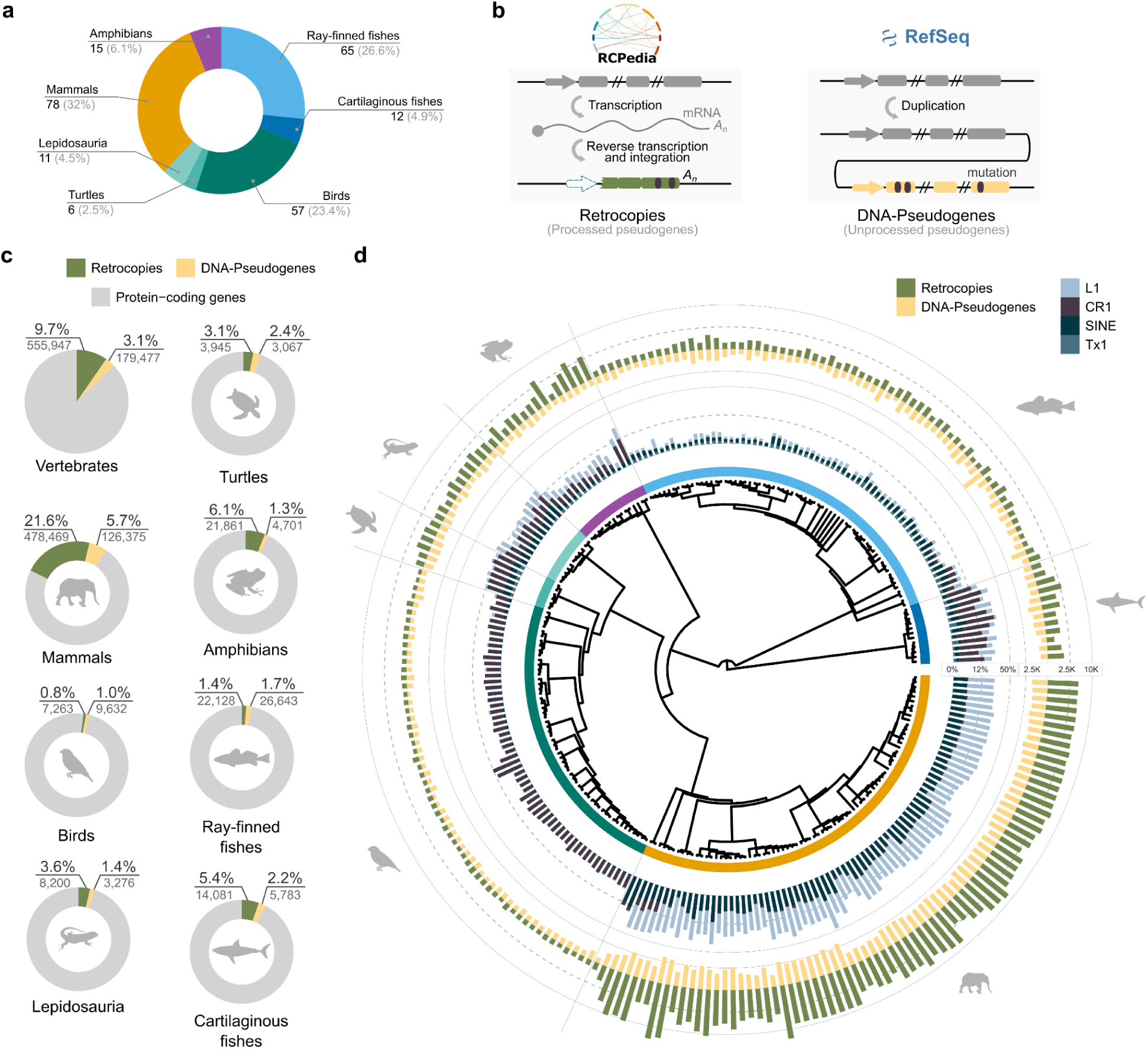
| Identification and profile of retrocopies and DNA-pseudogenes across vertebrate genomes. **a,** Taxonomic distribution of the 244 vertebrate species analyzed in this study. **b,** Schematic representation of the distinct molecular mechanisms generating retrocopies and DNA-pseudogenes, as well as the methodologies used for their identification. **c,** Genomic composition across vertebrate lineages showing the relative proportions of protein-coding genes (grey), retrocopies (green), and DNA-pseudogenes (yellow). Percentages and absolute counts are indicated for each class. **d,** Outer ring: per-species retrocopy (green) and DNA-pseudogene (yellow) counts (log₁₀ scale) across the vertebrate phylogeny. Inner ring: genomic proportions of LINE/L1 (light blue), LINE/CR1 (brown), LINE/Tx1 (blue), and SINE (dark blue) elements.

Mapping these counts onto the vertebrate phylogeny revealed pronounced lineage-specific patterns (**Fig. 1d**, outer ring; **Extended Data Fig. 1b**; **Supplementary Figs. 2-6**; **Supplementary Table 2**). Among mammals, Xenarthra (sloths in particular) reached ∼16,000 retrocopies per genome, consistent with a lineage-specific burst approximately 30 Mya^16^, while primates maintained broadly stable repertoires (∼8,000 in great apes)^3^ with expansions in specific taxa (*Nycticebus coucang*, ∼15,000) (**Extended Data Fig. 1b**; **Supplementary Table 2**). Rodentia, Chiroptera, Carnivora, and Cetacea showed intermediate but variable burdens (**Supplementary Fig. 2; Supplementary Table 2**). Outside mammals, retrocopy-dominated profiles in amphibians and lepidosaurs contrasted with the uniformly reduced repertoires of birds, conserved across deep avian divergences (**Extended Data Fig. 1b**; **Supplementary Figs. 3-6**; **Supplementary Table 2**).

Retrocopy abundance closely tracked autonomous non-LTR retrotransposon content (**Fig. 1d**, inner ring; **Supplementary Note 1**). In mammals, high LINE/L1 content was associated with elevated retrocopy counts, with Short Interspersed Nuclear Element (SINE) enrichment further reflecting *trans* utilization of L1 reverse-transcriptase machinery^17^ (**Extended Data Fig. 2a-c**). In non-mammalian lineages, LINE/CR1 or LINE/Tx1 elements instead sustained retroduplication (Extended Data Fig. 2a-c). Across species, retrocopy counts correlated strongly with the genomic fraction of autonomous non-LTR elements (Spearman ρ = 0.68, P = 3.8 × 10^-27^; **Extended Data Fig. 2d**; **Supplementary Fig. 7**). This relationship remained significant under phylogenetic generalised least squares (PGLS; β = 0.036, P = 0.0097) but explained only a fraction of the variance (R² ∼ 0.023; λ ∼ 0.97; **Extended Data Fig. 2f**), indicating that lineage-specific factors (e.g., germline transcriptome composition and chromatin accessibility^4^) outweigh current non-LTR retrotransposon content as determinants of retrocopy abundance. Some amphibians approach mammal-like retrocopy counts despite distinct retrotransposon profiles; conversely, lineages with depleted non-LTR content (ray-finned fishes and birds) consistently exhibit low counts (**Fig 1d**; **Extended Data Fig. 2a-d**).

### Conserved formation bias and lineage-specific sculpting shape retrocopy repertoires

We next examined how retrocopy repertoires partition between conserved and species-specific copies (**Fig. 2a**; **Supplementary Table 3**; **Supplementary Fig. 8**). Mammals (54.3% conserved), birds (48.7%), and turtles (52.6%) retained roughly balanced partitions, whereas amphibians (98.6% species-specific), cartilaginous fishes (95.7%), and lepidosaurs (90.6%) were overwhelmingly dominated by species-specific retrocopies, whereas ray-finned fishes displayed an intermediate pattern (71.7% species-specific). Globally, only 0.4% of retrocopies (1,932) were shared across distinct lineages (**Extended Data Fig. 3a-b**); the vast majority were restricted to within-order species (**Fig. 2b**; **Extended Data Fig. 3c**; **Supplementary Table 4**), consistent with a retroduplication pattern previously described in primates^3^ and here generalized across vertebrate diversity. Synonymous divergence (dS) between retrocopies and their parental genes independently corroborated these patterns (**Fig. 2c**): lineages with active retrotransposition (mammals, amphibians) showed sharp peaks at low dS values, consistent with abundant recent retroduplication, while lineages with reduced activity (e.g., birds and ray-finned fishes) exhibited flatter, right-shifted profiles matching their higher cross-order conservation fractions.

**Fig. 2.**
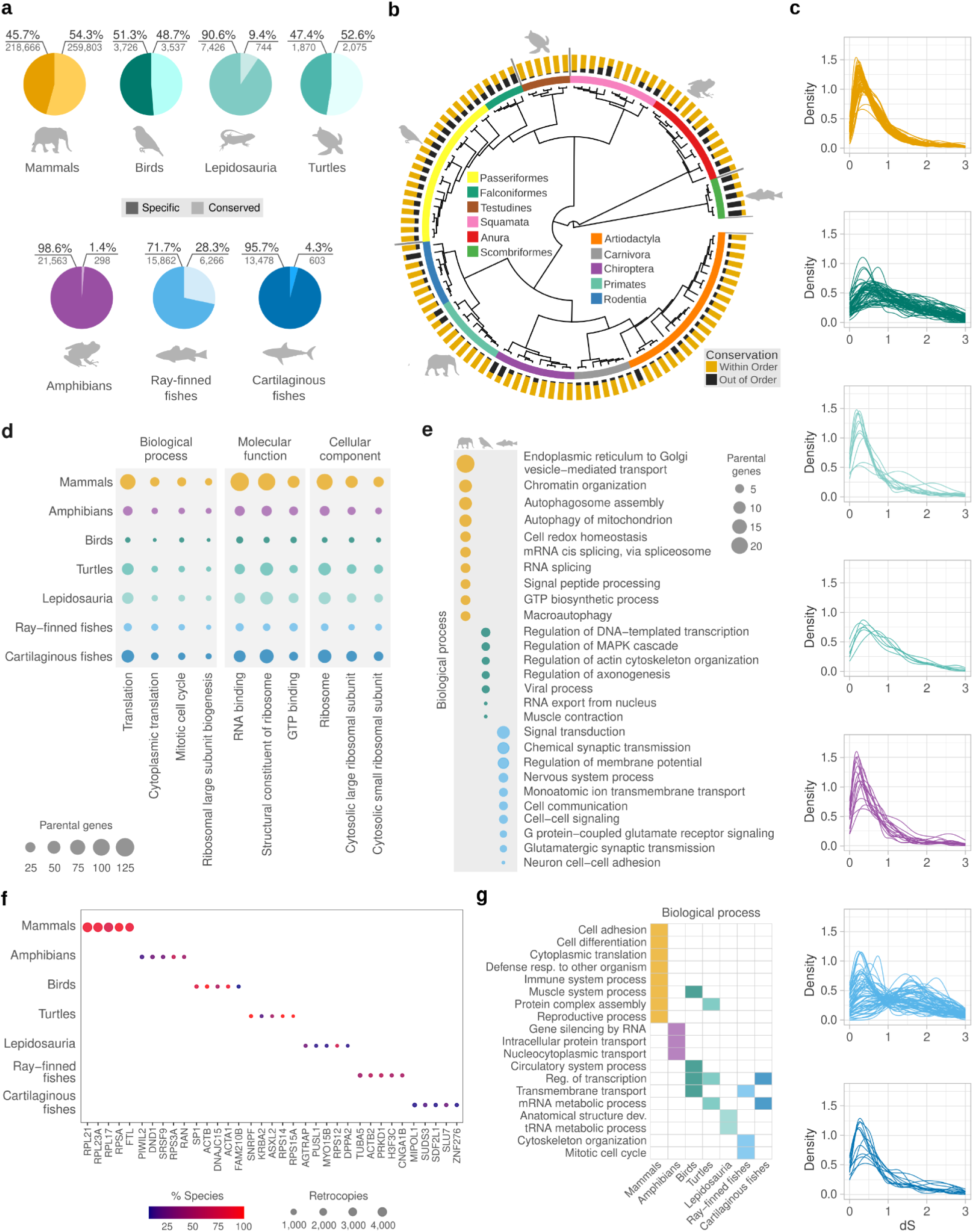
| Conserved formation bias and lineage-specific sculpting shape retrocopy repertoires across vertebrates. **a,** Proportion of species-specific (dark) and conserved (light) retrocopies for each major vertebrate lineage. **b,** Phylogenetic depth of retrocopy conservation for taxonomic orders. For each species, the proportion of conserved retrocopies shared within the same order (gold) versus across different orders (black) is shown (orders with more than five sampled species were used). **c,** Distributions of synonymous divergence (dS) between retrocopies and their parental genes, used as a proxy for retrocopy age. Each line represents one species. **d,** Gene Ontology (GO) terms significantly enriched (adjusted P < 0.05) among retrocopy parental genes across all seven vertebrate lineages. **e,** Lineage-restricted GO Biological Process enrichments (adjusted P < 0.05). Only lineages with significant lineage-specific enrichments are shown. **f,** Top five most frequently retrocopied parental genes within each vertebrate lineage. **g,** GO (Biological Process) annotations of the top retrocopied parental genes shown in f.

Despite this variation in conservation depth, Gene Ontology (GO) enrichment of retrocopy parental genes revealed a core set of functions significantly enriched across all seven lineages, centred on translation, RNA binding, structural constituent of the ribosome, and cytosolic ribosomal subunits (**Fig. 2d**; **Extended Data Fig. 4**; **Supplementary Table 5**). Relaxing the threshold to terms shared by six or five lineages expanded this core to include mRNA splicing, translational elongation, mitochondrial translation, and protein refolding (**Extended Data Fig. 4**). The consistency of this housekeeping signature across more than 500 million years of vertebrate divergence indicates that retroduplication repeatedly and independently captures transcripts from highly expressed genes involved in certain specific functions — a convergent process driven by transcriptional availability in the germline rather than shared ancestry of the retrocopies themselves.

Lineage-restricted enrichments, however, reveal that the functional composition of retained retrocopy repertoires is not fully explained by formation opportunity (**Fig. 2e**; **Extended Data Fig. 5a**). Mammals displayed the broadest spectrum, including chromatin organisation, autophagy, and MAPK cascade regulation. Ray-finned fishes showed a concentration of neural and synaptic processes (chemical synaptic transmission, membrane potential regulation, glutamatergic signalling). Birds, despite low retrocopy burdens, contributed enrichments in axonogenesis regulation and muscle contraction, indicating that even in TE-depleted genomes, the retained retrocopies derive from functionally non-random genes. Within mammals, order-level analysis revealed additional heterogeneity layered on the shared translational core, with several order-restricted biological processes among retrocopy parental genes (e.g., protein maturation in Artiodactyla and muscle system process in Primates). Notably, the monotreme profile was the most divergent overall, consistent with its basal phylogenetic position. (**Extended Data Fig. 5b**; **Supplementary Table 6**). A parallel dichotomy was observed when directly comparing parental genes of conserved versus species-specific retrocopies: conserved copies derived from core cellular processes (translational initiation, mitochondrial assembly, ubiquitin-dependent catabolism), while species-specific copies derived from more specialized regulatory programs (**Extended Data Fig. 5c**; **Supplementary Table 7**).

The most frequently retrocopied parental genes and their respective GO biological processes reinforced this dual pattern (**Fig. 2f-g**; **Supplementary Tables 8-11**). In mammals, four of the five top parental genes encode ribosomal proteins (*RPL21*, *RPL23A*, *RPL17*, *RPSA*), joined by *FTL*, which are all constitutively expressed transcripts, confirming that transcriptional availability drives retrocopy formation in L1-rich genomes (**Extended Data Fig. 6a-b**). Non-mammalian lineages diverged from this ribosomal core (**Fig. 2f-g**). In amphibians, the top parental genes include *PIWIL2* and *DND1* (germline piRNA and germ cell regulators) alongside *SRSF9* and *RAN*; the prominence of *PIWIL2*, a primary suppressor of transposon mobilization, is notable because it suggests a feedback dynamic in which the piRNA defence system is itself amplified by the retrotransposition machinery it restrains^18^. Birds showed a fully divergent profile (*SP1*, *ACTB*, *ACTA1*, *DNAJC15*, *FAM210B*), with no ribosomal proteins among the top five, consistent with post-formation selective retention of a narrow functional subset. Ray-finned fishes were dominated by cytoskeletal and signalling genes’ retrocopies (*TUBA5*, *ACTB2*, *PRKD1*, *H3F3C*, *CNGA1B*), and cartilaginous fishes by transcriptional regulators and stress-response genes’ retrocopies (*MIPOL1*, *SUDS3*, *SDF2L1*, *SLU7*, *ZNF276*).

Together, these results support a two-stage model of retrocopy evolution. In the first stage, a conserved transcription-driven formation bias repeatedly captures housekeeping transcripts across independent lineages, generating a shared substrate pool determined by germline expression levels. In the second stage, lineage-specific selective filtering sculpts the retained repertoire into distinct functional profiles — a process increasingly prominent in lineages with lower retrotransposon activity, where the small retrocopy complement that persists is enriched for functionally non-random categories. The analyses that follow provide multiple lines of evidence supporting this model across three domains: selective constraint and chromosomal redistribution, species life-history traits, and cancer gene retroduplication in relation to body size and longevity.

### ORF retention and selective constraint identify functionally recruited retrocopies

All lineages harboured retrocopies retaining intact open reading frames (**Fig. 3a**; **Supplementary Table 12**). Amphibians and mammals showed the highest ORF-bearing fractions (34.0% and 33.1%, respectively), consistent with their elevated recent retrotransposition and younger age profiles. Birds (27.1%), turtles (23.1%), ray-finned fishes (23.2%), and lepidosaurs (22.5%) occupied intermediate positions, while cartilaginous fishes exhibited the lowest ORF fraction (15.7%).

**Fig. 3.**
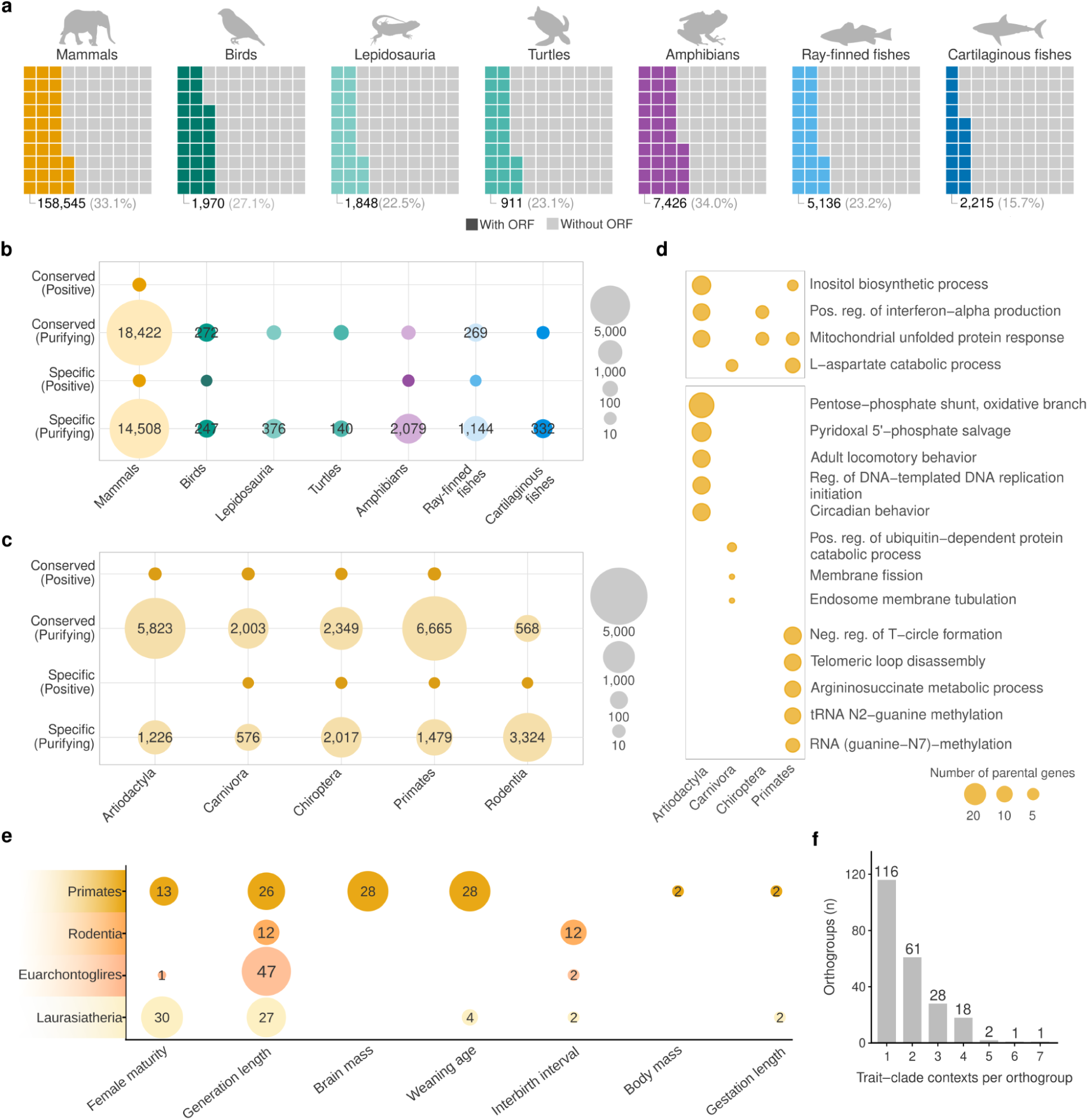
| ORF retention, selective constraint, and life-history associations shape the retrocopy repertoire across vertebrates. **a,** Fraction of retrocopies retaining an intact ORF in each vertebrate lineage. Each square represents 1% of the lineage-specific retrocopy complement; absolute counts and percentages are indicated below. **b,** Numbers of ORF-bearing retrocopies evolving under purifying or positive selection, stratified by conservation status (conserved vs. species-specific). Dot size reflects retrocopy count. **c,** As in b, resolved across major mammalian orders. **d,** GO Biological Process enrichment among parental genes of constrained retrocopies within mammalian orders. Upper panel: terms enriched across at least two mammalian orders; lower panel: order-specific enrichments. Dot size reflects the number of parental genes. **e,** Number of orthogroups with significant associations (based on phylogenetic linear regressions) between retrocopy fraction and life-history traits across four mammalian orders. Circle size and number indicate the count of significantly associated orthogroups. **f,** Distribution of orthogroups by the number of traits–orders contexts in which they are significant.

To distinguish ORF retention due to young age from genuine functional constraint, we applied dN/dS-based tests, where dN and dS represent the rates of non-synonymous and synonymous substitutions, respectively, classifying ORF-bearing retrocopies as evolving under neutral, purifying, or positive selection and stratifying by conservation status (**Fig. 3b**; **Supplementary Fig. 9**; **Supplementary Table 12**). Purifying selection predominated across all lineages. Mammals contributed the largest constrained set, with 18,422 conserved and 14,508 species-specific retrocopies under purifying selection. Substantial constrained sets were also detected in amphibians (2,079 species-specific) and ray-finned fishes (269 conserved; 1,144 species-specific). Candidates for positive selection were rare (primarily mammals: 41; all others combined: 6), indicating that long-term coding retrocopy retention is dominated by purifying constraints rather than pervasive adaptive protein evolution.

Within mammals, the constrained compartment showed pronounced order-level structure (**Fig. 3c**). Primates and Artiodactyla carried the largest conserved constrained sets (6,665 and 5,823 under purifying selection, respectively), Rodentia showed the largest species-specific compartment (3,324) consistent with elevated recent turnover, and Chiroptera displayed a balanced profile (2,349 conserved; 2,017 species-specific), suggesting both sustained ancient retention and ongoing recruitment. GO annotation of parental genes giving rise to order-restricted constrained retrocopies revealed distinct functional signatures for each order (**Fig. 3d**; **Supplementary Table 13**). Artiodactyla displayed the broadest profile, spanning inositol biosynthesis, mitochondrial unfolded protein response, interferon-alpha regulation, and circadian behaviour — predominantly driven by Cetacea species and consistent with metabolic adaptations linked to the aquatic transition^19^. Chiroptera shared two of these processes (interferon-alpha production and mitochondrial unfolded protein response), a convergence notable given that both orders harbour lineages with distinctive immune adaptations: antiviral tolerance in bats^20^ and aquatic immune challenges in Cetacea^21^. Primates were dominated by telomere biology^22^ and RNA modification^23^, pointing to selective retention of retrocopies involved in genome stability and translational regulation. These order-specific and partially convergent functional biases provide direct evidence for the second stage of the model: lineage-specific selective filtering shapes the constrained retrocopy repertoire according to order-specific physiological demands, while shared processes (particularly innate immunity and mitochondrial homeostasis) indicate convergent selective pressures acting independently on retrocopy retention across mammalian lineages.

Beyond molecular signatures, retrocopy dosage variation correlates with mammalian life-history traits at the phenotypic level. Phylogenetic regressions identified 84 significant associations between orthogroup-level retrocopy fraction and traits including brain mass, generation length, and reproductive timing, the majority confined to single trait-lineage contexts — as predicted by the two-stage model (**Fig. 3e-f**; **Supplementary Table 14**). In primates, orthogroups linked to brain mass were enriched for glucose import and carbohydrate transport, consistent with the energetic demands of neurodevelopment (**Extended Data Fig. 7b**).

### Sex chromosome-autosome retrocopy trafficking reveals MSCI-driven asymmetry in mammals

mRNA retroduplication provides a mechanism by which genes can escape chromosomal regulatory constraints, most notably meiotic sex chromosome inactivation (MSCI), the transcriptional silencing of X and Y chromosomes during male meiotic pachytene^24,25^. Autosomal retrocopies of X-linked genes have been proposed to compensate for this silencing during spermatogenesis^26^. To test this model across vertebrate diversity, we mapped the chromosomal origin and destination of all retrocopies in species with XY (mammals) and ZW (birds and reptiles) sex-determination systems (**Fig. 4a-d**; **Supplementary Tables 15-16**).

**Fig. 4.**
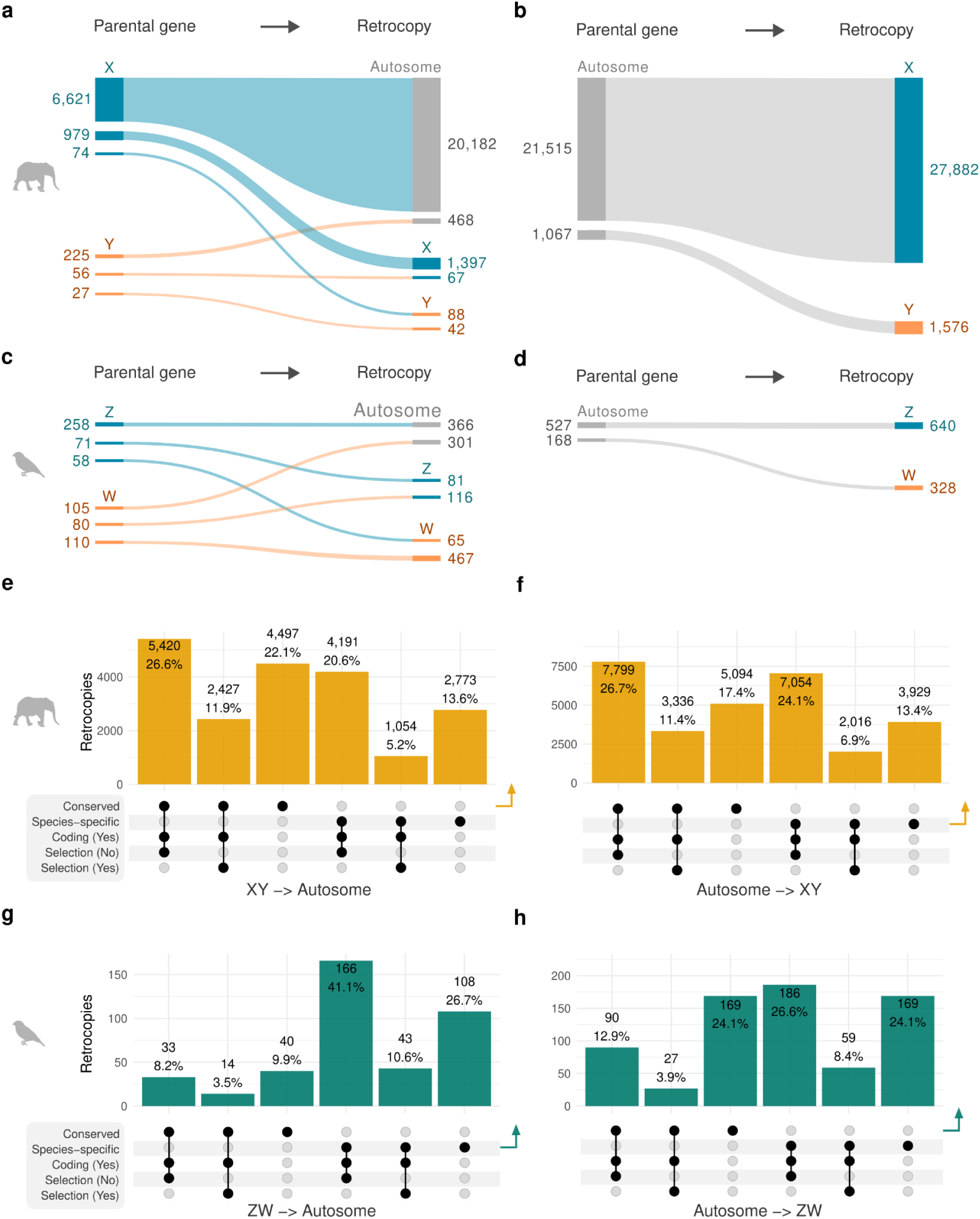
| Sex chromosome–autosome retrocopy trafficking and functional asymmetry in mammals and birds. **a-b,** Retrocopy flux between sex chromosomes (X, Y) and autosomes in mammals. **a,** parental genes on sex chromosomes and the chromosomal destinations of their retrocopies. **b,** autosomal parental genes producing retrocopies on the X or Y. Numbers on the left indicate parental gene counts; numbers on the right indicate retrocopy counts. **c-d,** equivalent a-b analysis for the ZW sex-determination system (birds and reptiles). **e-f,** Upset intersection plots stratifying mammalian sex-autosome retrocopies by conservation status, coding potential, and purifying selection signatures. **e,** XY-to-autosome retrocopies; **f,** autosome-to-XY retrocopies. Arrows highlight the most functionally constrained category (conserved, coding, under selection). Percentages indicate the proportion of retrocopies in each category relative to the total for that trafficking direction. **g-h,** as in **e-f**, for avian ZW-autosome trafficking. Bar colours follow the lineage colour scheme used throughout (gold, mammals; teal, birds).

In the mammalian XY system, the dominant retrocopy flux originated from the X chromosome (**Fig. 4a-b**): 6,621 X-linked parental genes generated 20,182 autosomal retrocopies, yielding a retrocopy-per-parental ratio of 3.05 — significantly above the genomic average of 2.87 (P = 2.89x10⁻⁵; Chi-square: 17.48; **Supplementary Table 17**). Of all retrocopies generated by X-linked genes, 93.1% landed on autosomes, with only 6.4% remaining on the X and 0.4% reaching the Y (**Fig. 4a**). This pronounced export asymmetry was unique to the X: Y-linked genes showed a lower ratio (2.08) but still significantly above the genomic average (P = 6.78x10⁻⁵; Chi-square: 15.87), whereas cross-traffic between sex chromosomes was rare (**Fig. 4a**).

In the avian ZW system, where MSCI does not operate, the pattern was diverged from the mammalian one (**Fig. 4c-d**). Z-linked genes showed a retrocopy-per-parental ratio significantly below the genomic average (1.42 vs. 1.71; P = 0.019), and only 65.0% of Z-derived retrocopies landed on autosomes — a markedly weaker export signal than the 93.1% for the mammalian X chromosome (**Fig. 4a**). This absence of a strong “out of the Z” pattern provides a natural control: in ZW species the directional bias characteristic of the mammalian X does not occur. The W chromosome displayed a distinct profile: W-linked genes generated retrocopies at a rate above the genomic background (2.87 vs. 1.71; P = 3.2 × 10⁻⁶), but unlike the mammalian X most did not escape the W — 56.1% remained on the W itself, consistent with the low recombination and reduced purifying selection on a degenerating chromosome^27^.

The out-of-X signal extended beyond retrocopy volume to the evolutionary properties of exported copies (**Fig. 4e-h**). In mammals, retrocopies moving from sex chromosomes to autosomes were enriched (P = 5.1 x 10^-9^; chi-square: 34.14) for conserved, coding sequences or under purifying selection, comprising 60.6% of the XY-to-autosome retrocopy flux (**Fig. 4e**). Retrocopies in the reverse direction (autosomes to sex chromosomes) were instead enriched for species-specific insertions (**Fig. 4f**). In birds, the pattern was inverted: the ZW-to-autosome traffic was dominated (P = 0.0054; chi-square: 7.75) by species-specific copies (84.4%, **Fig. 4g**), whereas the autosome-to-ZW flux showed a higher proportion of constrained retrocopies (**Fig. 4h**). This reciprocal asymmetry constitutes a natural quasi-experiment: in mammals, where MSCI silences the X during spermatogenesis, functional retrocopies are preferentially retained when they escape to autosomes; in ZW species, no such directional enrichment occurs — instead, functional copies are associated with movement into the sex chromosomes.

The breadth of this pattern is exemplified by canonical cases. *PGK2* and *PDHA2*, autosomal retrocopies of X-linked glycolytic genes *PGK1* and *PDHA1*, compensate for MSCI-mediated silencing in testis^28,29^. Our data show that coding retrocopies of *PGK1* and *PDHA1* are under purifying selection in 36 and 29 mammalian species, respectively, extending these examples from model organisms to a pan-mammalian pattern (**Supplementary Tables 18-19**). Beyond these classical cases, SMS (spermine synthase) emerges as a recurrent out-of-X target, with independently generated autosomal retrocopies retaining coding potential and purifying selection signatures across Carnivora, Chiroptera, Rodentia, and Primates — orders that diverged ∼90-100 Mya^30^ (**Extended Data Fig. 8a-b**; **Supplementary Table 20**). The functional relevance of SMS retroposition is supported by its essential role in polyamine biosynthesis for sperm viability^31^ and its association with Snyder-Robinson syndrome when mutated^32^.

### Cancer gene retrocopies reveal tumor suppressor bias in large-bodied and long-lived lineages

Cancer-associated genes are of particular interest as retrocopy substrates because most are highly expressed in the germline and their copy-number variations have established consequences for cancer susceptibility^33–35^. We classified cancer genes based on human annotations^36^ as tumor suppressor genes (TSGs; 66 genes) or oncogenes (41 genes), **Supplementary Table 21**. Both classes showed a striking propensity for retrotransposition: 81.8% of TSGs (54/66) and 80.5% of oncogenes (33/41) had at least one retrocopy in at least one VGP species, proportions significantly exceeding the genome-wide baseline (**Fig. 5a**; P = 4.08 × 10^-20^, Fisher’s exact test; **Supplementary Table 22**).

**Fig. 5.**
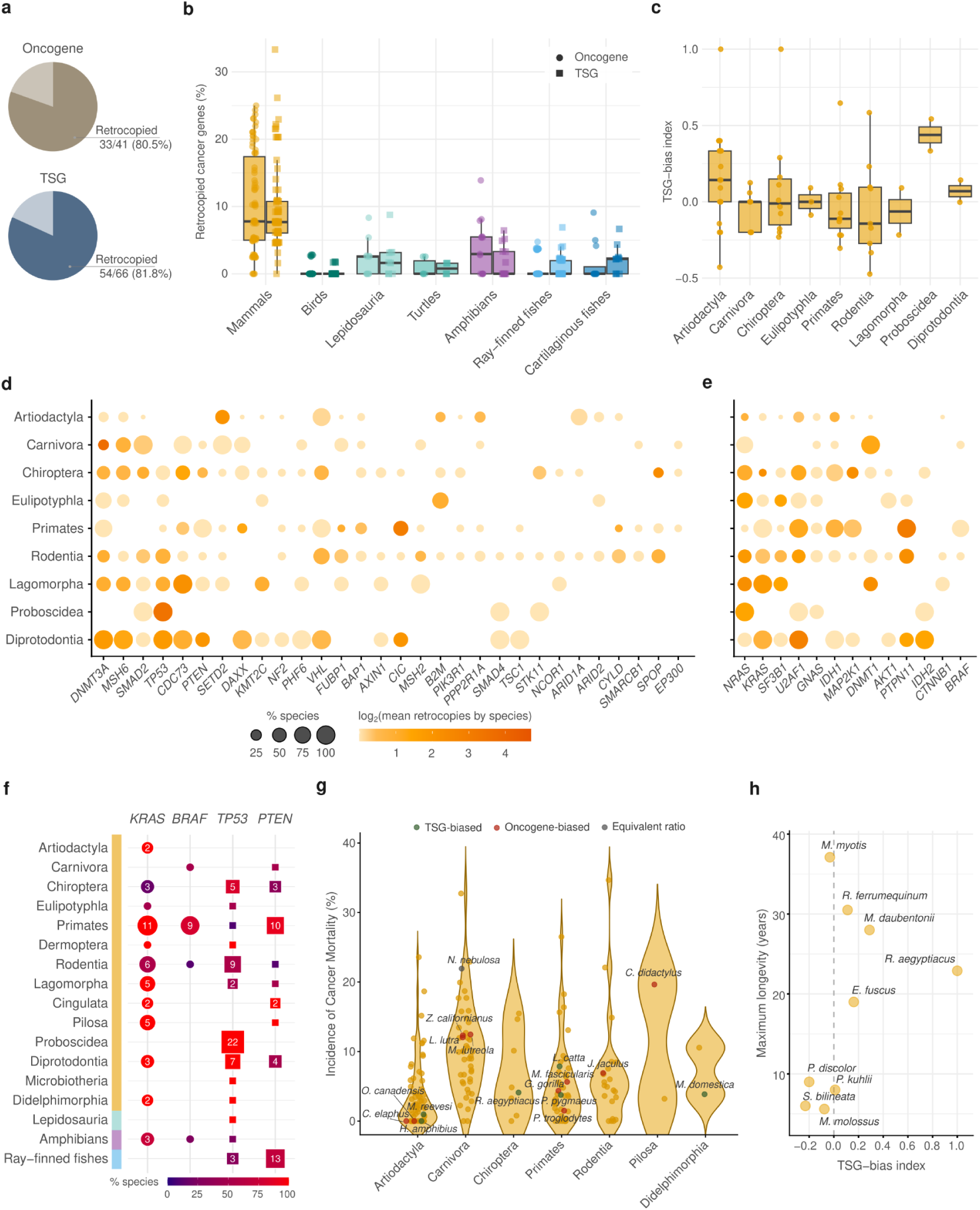
| Retrocopies of cancer-associated genes across vertebrate genomes reveal a tumor suppressor bias in large-bodied and long-lived lineages. **a,** Proportion of tumor suppressor genes and oncogenes that have generated at least one retrocopy in at least one VGP species. **b,** Per-species percentage of retrocopied cancer genes grouped by vertebrate lineage. Individual species are overlaid as points (TSGs, squares; oncogenes, circles). **c,** TSG-bias index — defined as (TSG − oncogene) / (TSG + oncogene) retrocopies — per species, grouped by mammalian order. Values above zero indicate preferential accumulation of TSG retrocopies; values below zero indicate oncogene bias. **d,** Retrocopy profile of individual TSGs across nine mammalian orders. Dot size reflects the percentage of species within the order harbouring at least one retrocopy of the indicated gene; colour intensity reflects log₂(mean retrocopy count per species). **e,** As in d, for oncogenes. **f,** Retrocopy counts and species prevalence for four focal cancer genes: *KRAS* and *BRAF* (oncogenes) and *TP53* and *PTEN* (TSGs), across mammalian orders and non-mammalian vertebrate lineages. Dot size reflects the percentage of species with retrocopies; numbers indicate total retrocopy counts. Background colour bars denote lineage. **g,** Incidence of cancer mortality (%) for VGP species with available zoo-based data, grouped by mammalian order. Point colour indicates TSG-bias category: green, TSG-biased; red, oncogene-biased; gray, equivalent ratio; yellow, species still out of VGP. **h,** Association between maximum longevity and the TSG-bias index in Chiroptera.

Across vertebrate lineages, cancer gene retrocopy content mirrored the overall retrocopy landscape (**Fig. 5b)**: mammals harboured the highest per-species proportions (P < 10^-16^; two-sided Wilcoxon rank-sum test), with individual species reaching over 30% of TSGs retrocopied (**Extended Data Fig. 9a**); birds and ray-finned fishes showed the lowest rates (median <1%; **Fig 5b**). The substantial mammalian retrocopy burden, combined with sufficient interspecific variation, motivated us to focus subsequent analyses on this lineage. To quantify the balance between TSG and oncogene retrocopies, we computed a TSG-bias index per species, defined as (TSG − oncogene) / (TSG + oncogene) retrocopies (**Supplementary Table 23**). Orders associated with low cancer mortality, including Proboscidea ^37^ and Artiodactyla^38^, displayed consistently positive values indicating preferential TSG retrocopy accumulation, whereas Rodentia, Primates, and Lagomorpha showed negative values reflecting relative oncogene enrichment (**Fig. 5c**).

Analysis of individual cancer genes across mammalian orders revealed a structural asymmetry (**Fig. 5d-e**; **Supplementary Table 24**). TSGs displayed a broad retrocopy landscape: 55.5% (30/54) harboured retrocopies in three or more orders, compared to only 39.3% (13/33) of oncogenes. TSG retrocopies were distributed across a wide gene repertoire — *PTEN*, *TP53*, *CDC73*, *SMAD2*, and *CIC* each appearing independently in five or more orders — whereas oncogene retrocopies concentrated in fewer genes (*NRAS*, *U2AF1*, *IDH1*) and fewer orders (**Fig. 5d-e)**. This asymmetry is consistent with differential post-insertion retention: additional tumor suppressor copies are selectively neutral or beneficial, whereas ectopic oncogene expression may carry deleterious gain-of-function effects. Notably, even in orders with large retrocopy complements across both panels (Chiroptera, Rodentia), the TSG repertoire remained substantially wider, indicating that the asymmetry reflects differential retention rather than formation bias — a direct manifestation of stage two of the two-stage model.

Four genes with well-characterized retrocopies illustrate the evolutionary dynamics (**Fig. 5f**; **Supplementary Table 24**). *TP53* showed the most expansive signal (**Fig. 5f**; **Extended Data Fig. 9b**): 22 retrocopies in Proboscidea, 9 in Rodentia, 7 in Diprotodontia, 5 in Chiroptera, with additional copies in amphibians, lepidosauria, and ray-finned fishes, extending the footprint of *TP53* retrotransposition well beyond mammals^39,40^. *PTEN* displayed fewer expansion across lineages, however with high copy numbers (10 in Primates, 13 in ray-finned fishes, and other 13 across 7 mammals orders; **Fig. 5f; Extended Data Fig. 9c**), consistent with intrinsic transcript features driving recurrent retrotransposition and the known ceRNA function of *PTENP1*^41^. Among oncogenes, *KRAS* was the most taxonomically widespread retrocopy (**Fig. 5f; Extended Data Fig. 9d**), present across 11 mammalian orders and in 3 amphibian species, yet its ubiquity contrasts with the narrower overall oncogene landscape — reinforcing that retrocopy formation is not impeded for oncogenes but selective retention is restricted. *BRAF* retrocopies were the most taxonomically confined (Primates, Rodentia, Carnivora, and one amphibian species; **Extended Data Fig. 9e**). Notably, a Cetacea-specific retroduplication of *CDKN2C* in bowhead whales (**Supplementary Table 24)** has been implicated in longevity and tumor suppression through promoter cooption and constitutive tissue expression^42^, providing a mechanistic precedent for functional incorporation of TSG retrogenes into cancer-resistance programmes.

We then asked whether the TSG-bias index correlates with cancer-relevant phenotypes. Cross-referencing VGP species with zoo-based cancer mortality data^38,43^, species with positive TSG-bias tended to cluster at lower cancer mortality, whereas oncogene-biased species showed more variable rates (**Fig. 5g**). Moreover, within Chiroptera — an order notable for exceptional longevity relative to body size and low cancer rates — we observed a positive association between maximum lifespan and TSG-bias index (**Fig. 5h**; **Supplementary Table 25**): three of four species with longevity exceeding 10 years exhibited TSG-biased profiles (*R. ferrumequinum*, *M. daubentonii, R. aegyptiacus, and E. fuscus*), while all four species below 10 years were oncogene-biased, although this pattern is based on limited sampling (8 species). At a broader scale, large-bodied mammals (>70 kg) exhibited significantly higher TSG-bias indices than small-bodied species (P = 0.039, two-sided Wilcoxon rank-sum test; **Extended Data Fig. 9f**), with the most prominent small-bodied TSG-biased outliers being bats — including *Rousettus aegyptiacus*, a species with an exceptionally long lifespan for its body mass.

Together, these findings reveal that cancer gene retrotransposition is a recurrent and evolutionarily structured process in vertebrates, especially in mammals. The positive association between TSG retrocopy bias, longevity, and body size identifies retrocopy-mediated expansion of tumor suppressor dosage as a previously unrecognized genomic correlate of Peto’s paradox^44^ (**Supplementary Note 2**), and establishes retroduplication as an underappreciated mechanism by which vertebrate genomes have repeatedly expanded their tumor-suppressive capacity.

### A comprehensive retrocopy resource for the vertebrate genomics community

To provide a publicly accessible platform for investigating retrocopy biology across vertebrate diversity, as well as exploration and extension of these findings, we integrated the complete retrocopy catalogue into the RCPedia database (https://www.rcpedia.org/vgp), providing species-level annotations for all 244 VGP genomes — including parental gene assignments, genomic coordinates, conservation status, ORF predictions, and dN/dS-based selection classifications. All annotations are additionally available as a dedicated track in the UCSC Genome Browser (genome.ucsc.edu), where individual retrocopies can be visualized alongside gene models, conservation scores, and transposable element annotations.

## DISCUSSION

Our analysis of 244 chromosome-scale vertebrate genomes establishes retrocopies as dynamic, functionally consequential components of genome evolution rather than inert relics of retrotransposition. The two-stage model we establish — conserved formation bias followed by lineage-specific selective filtering — provides a framework that explains how identical retrotransposition machinery yields fundamentally different functional outcomes across vertebrate diversity. The formation stage is remarkably conserved: parental genes across all seven lineages converge on the same housekeeping functions, reflecting a universal germline-expression constraint on which transcripts are available for retroduplication. The filtering stage diverges: neural processes accumulate in ray-finned fishes, chromatin and secretory pathways in mammals, and axonogenesis in birds. This divergence intensifies in non-LTR retrotransposons depleted lineages (e.g., birds and ray-finned fishes), where the small retrocopy complement is disproportionately enriched for specific functional categories — precisely the pattern expected if post-insertion selection, rather than formation opportunity, shapes the retained repertoire.

Two biological domains provide independent lines of evidence supporting this two-stage model and illustrate its explanatory reach. Sex chromosome-autosome retrocopy trafficking reveals that the mammalian X chromosome exports retrocopies to autosomes at elevated rates (93.1%), enriched for conserved, coding sequences under purifying selection — extending the “out of the X” hypothesis^24,26^ mediated by mRNA retroduplication to unprecedented phylogenetic breadth. The avian ZW system provides a critical natural control: Z-linked genes show no equivalent export enrichment, and functional retrocopies are instead associated with the autosome-to-sex-chromosome direction. This reciprocal asymmetry — predicted by the presence of MSCI in mammals and its absence in birds — constitutes among the strongest comparative evidence that meiotic sex chromosome inactivation drives the directional redistribution of gene function via retroduplication.

Cancer gene retroduplication reveals a second axis of the model’s selective filtering. More than 80% of both TSGs and oncogenes have generated retrocopies, yet their post-insertion fates diverge: TSG retrocopies accumulate preferentially in large-bodied, long-lived lineages, while oncogene retrocopies are more taxonomically restricted. The convergent *TP53* expansion in Proboscidea, the positive association between TSG-bias and longevity in Chiroptera, and the significant TSG enrichment in larger mammals collectively position retroduplication as a contributor to resolving Peto’s paradox^44^ — adding retrocopy-mediated dosage expansion to the established mechanisms of gene duplication and positive selection^37^.

Several limitations warrant consideration. Conservation estimates remain sensitive to taxonomic sampling density. The cancer mortality data derive from zoo-based surveys^38,43^ that may not fully reflect wild incidence. The within-order correlations between TSG-bias and longevity — while suggestive — rest on small sample sizes (8 species) that limit statistical power, and the TSG-bias index may be influenced by the unequal number of TSGs (66) versus oncogenes (41) in the reference set. More broadly, the absence of matched multi-OMICs data across VGP species means that our functional inferences rest on ORF retention and selection signatures rather than direct evidence of expression or translation.

Future integration of transcriptomic, epigenomic, and population-level data will be essential for determining which retrocopies are actively expressed, translated, or incorporated into regulatory networks. The RCPedia database and UCSC Genome Browser track released with this study provide a foundation for such integrative efforts across vertebrate diversity.

In conclusion, retrocopies represent a continuously generated substrate for evolutionary innovation that each vertebrate lineage has independently shaped through the interplay of retrotransposon activity, germline transcription, and natural selection. The convergent patterns we describe — from the chromosomal redistribution of sex-linked genes to the expansion of tumor suppressor dosage in cancer-resistant lineages — reveal retroduplication as a pervasive yet overlooked axis of vertebrate genome evolution.

## METHODOLOGY

### Genome dataset

We analyzed genome assemblies from 244 vertebrate species, including mammals, birds, lepidosaurs, turtles, amphibians, and fishes, generated by the Vertebrate Genome Project^12^. Only assemblies with transcript annotations available through RefSeq were included (**Supplementary Table 1**).

### Identification of retrocopies

We identified retrocopies using an approach previously described and implemented by RCPedia^13^. This strategy consists of aligning multi-exonic protein-coding gene transcripts to the corresponding reference genome in order to detect intronless genomic regions.

Messenger RNA (mRNA) sequences were retrieved from genomic coordinates provided by RefSeq for each assembly using gffread^45^ (v0.12.7; default parameters). The extracted mRNA sequences were aligned against their respective reference genomes using LAST^46^ (v1066; parameters: lastal-D1000) to identify candidate retrocopies. Alignments shorter than 120 bp were discarded.

To avoid detecting chromosomal duplications, candidate events located within 200,000 bp of the parental gene were excluded. Additionally, candidate retrocopies were required to preserve at least one of the three terminal exon-exon junctions of the parental gene. This filter was designed to reflect the biological mechanism of retrotranscription initiated from the poly(A) tail. Alignments containing ≥40% repetitive elements were removed. Repetitive regions were defined based on genomic annotations provided^12^, including lineage-specific interspersed repeat annotations generated by RepeatModeler and masked with RepeatMasker, simpleRepeats, and windowMasker. To prevent tandem duplication artifacts of the same retrocopy, events derived from the same parental gene were required to be separated by more than 500,000 bp. Candidate events overlapping more than three exons of annotated protein-coding genes were excluded. Likewise, parental genes presenting more than five retrocopies overlapping annotated protein-coding genes were discarded. These filters were implemented to reduce false positives arising from potential annotation artifacts.

In cases of multiple candidate parental genes, the parental assignment was resolved by selecting the gene with the highest match/(match + mismatch) ratio. Additionally, nearby alignments (≤6,000 bp apart) were merged to account for fragmented alignments potentially caused by repetitive elements.

### Classification of DNA-pseudogenes

DNA-pseudogene annotations were derived from the “pseudogenes” category in the RefSeq genome annotations. To classify annotated events as DNA-pseudogenes, a species-specific size threshold was applied. For each species, the median exon length and median intron length of protein-coding genes were calculated. The following formula was then used to define the minimum expected size of a DNA-pseudogene: (2 × median exon length) + median intron length. Annotated pseudogene events with lengths equal to or greater than this threshold were classified as DNA-pseudogenes. We also discarded pseudogenes that were classified as retrocopies in the previous step.

### Orthology inference of protein-coding genes

Orthologous relationships among protein-coding genes (from RefSeq annotations) were retrieved from the NCBI Gene database (https://ftp.ncbi.nlm.nih.gov/gene/DATA/gene_orthologs.gz). Orthology assignments were represented as an undirected graph using the R package igraph (v2.2.0)^47^, in which nodes correspond to gene-species pairs and edges denote orthologous relationships. Connected components in this graph were used to define orthologous gene clusters across species. Each gene was then annotated with its corresponding taxonomic lineage and order. For each lineage and order, we quantified the number of protein-coding genes whose orthologs were restricted to the same group (within-lineage/order conservation) or extended to other groups (across-lineage/order conservation), relative to the total number of protein-coding genes. To assess differences in conservation patterns between retrocopies and protein-coding genes, we compared the proportion of genes conserved across taxonomic groups using Fisher’s exact tests.

### Synteny-based orthology inference

To infer orthologous relationships among the detected retrocopies, we employed a synteny-based approach. For each retrocopy, the genomic sequence corresponding to the retrocopy locus plus 3,000 bp flanking regions on both sides was retrieved. Pairwise sequence alignments were performed using LASTZ (v1.04.29; --format=general [multiple]; https://lastz.github.io/lastz/). For downstream analyses, we retained alignments with sequence coverage ≥50% and sequence identity ≥60%. Additionally, we required that at least 50% of the retrocopy region itself be aligned.

In cases of multiple alignments for a given locus, the alignment with the highest coverage was selected. Based on Order classification available^12^, orthology relationships were classified into three categories: (1) Species-specific: retrocopies with no alignments meeting the defined criteria in other species; (2) Order-conserved: retrocopies conserved in at least one species belonging to the same taxonomic order; and (3) Out-of-order: retrocopies conserved in at least one species outside the taxonomic order of origin.

### dN/dS analysis

To estimate synonymous substitutions per synonymous site (dS) and nonsynonymous substitutions per non-synonymous site (dN), open reading frames (ORFs) were first identified in retrocopies using TransDecoder (v5.7.1; TransDecoder.LongOrfs -m 75)^48^, with a minimum ORF length of 75 amino acids. Only complete ORFs were retained for downstream analysis.

Predicted amino acid sequences for each retrocopy were aligned against the corresponding parental protein sequences obtained from RefSeq using ClustalW (v2.1; default parameters)^49^. Codon-aware nucleotide alignments were subsequently generated with PAL2NAL (v14; -output paml -nogap)^50^, based on the protein alignments and their corresponding coding sequences. Synonymous and nonsynonymous substitution rates were estimated using codeml from the PAML package^51^, with the following parameters: CodonFreq = 2, model = 0, Nsites = 0, fix_omega = 0, and omega = 0.4.

### Construction of mammalian orthogroup-by-species retrocopy count matrices

For each mammalian genome, we compiled a per-species mapping between parental genes and their associated retrocopy identifiers (RTCs) from the retrocopy annotation tables produced in our pipeline. In parallel, we parsed the NCBI genome feature tables for each species to extract explicit gene-transcript-protein relationships, thereby allowing parental genes to be unambiguously associated with protein identifiers. To assign orthology, we used the OrthoFinder orthogroup file (Orthogroups.txt) provided by the VGP consortium^12^. For each species, we linked parental genes to orthogroups through a two-step intersection: (i) parental gene identifiers were mapped to their corresponding protein IDs using the feature-table-derived gene-protein mapping, and (ii) these protein IDs were matched to OrthoFinder orthogroups based on their presence in the orthogroup file. For each species and orthogroup, we counted the number of distinct RTC records and pivoted the result to a wide matrix with species as rows and orthogroups as columns. To avoid extremely sparse orthogroups with insufficient phylogenetic breadth, we retained only orthogroups present (nonzero counts) in at least 20 mammalian species.

### Phylogenetic association tests with life-history traits

Life-history trait data, including body mass, lifespan, brain mass, female maturity, gestation length, litter size, interbirth interval, and weaning age, were obtained from a previously published dataset^52^.

Associations between orthogroup retrocopy fraction and life-history traits were tested using phylogenetic regression models, using the “phylolm” function in the R package phylolm (v2.6.5)^53^. For each trait, mammalian lineage subset, and orthogroup, retrocopy counts were analyzed against trait values using phylogenetic linear models, with trait preprocessing in log_10_ space. At least two alternative model structures were tested: (i) a Brownian-motion model (“BM”) and (ii) a Pagel’s λ model (“lambda”), returning effect sizes (slope), model fit statistics (AIC, R²), and an estimated lambda parameter, where applicable. The preferred model per orthogroup was selected by minimum AIC and the strength of model preference was summarized as ΔAIC between competing fits. Multiple-testing correction was applied within each trait versus mammalian-subset analysis (FDR; reported as q-values for BM and lambda). Only orthogroups with adjusted p-value ≤ 0.05 were used in downstream analysis.

### Definition of orthogroup sets for downstream overlap summaries and GO analysis

We defined “GO sets” as the collection of orthogroups significant for a given mammalian subset and trait (i.e., orthogroups grouped by subset vs. trait). Each set was assigned to a unique identifier set_id = subset_trait. The orthogroup universe for GO analyses was derived from the set of all orthogroups used for the phylogenetic association tests. To interpret significant sets functionally, we mapped orthogroups to human gene symbols. Because orthogroups can contain multiple human genes, this mapping is inherently many-to-many; we therefore defined the foreground gene list for each set_id as the union of unique human gene symbols assigned to orthogroups in that set. The GO background universe was defined as all human genes mapped from the orthogroup columns retained in the mammalian orthogroup count matrix (orthogroups present in ≥20 mammalian species). Human symbols were converted to Entrez IDs using bitr(org.Hs.eg.db)^54^, and GO Biological Process enrichment was performed with clusterProfiler::enrichGO() using the foreground Entrez IDs, the universe Entrez IDs, and Benjamini-Hochberg adjustment. We applied an FDR threshold (adjusted p-value ≤ 0.05) and optionally reduced redundancy among enriched GO terms using clusterProfiler::simplify() (Wang semantic similarity, cutoff = 0.7, retaining the most significant representative term). For visualisation and concise reporting, we extracted the top five enriched BP terms per set based on adjusted p-value.

### Functional annotation and enrichment analyses of retrocopied genes

Gene Ontology (GO) annotations for all analyzed species were obtained from the NCBI gene2go dataset (https://ftp.ncbi.nlm.nih.gov/gene/DATA/gene2go.gz). First, to summarise the functional landscape of retrocopied genes, GO full annotations were mapped to GO slim categories. The Gene Ontology structure was retrieved from the Gene Ontology Consortium^55^ (http://purl.obolibrary.org/obo/go.obo), and the generic GO slim ontology was obtained from http://current.geneontology.org/ontology/subsets/goslim_generic.obo. Ontologies were parsed using the ontologyIndex R package (v2.12)^56^, and mapping from full GO terms to GO slim categories was performed by traversing the Gene Ontology directed acyclic graph via is_a relationships. For each GO term, its full set of ancestral terms was identified, and any ancestors belonging to the GO slim subset were retained as valid mappings. GO slim term names and subontology assignments were retrieved using the GO.db package (v3.16.0)^57^. Redundant parental gene annotations within the same GO slim category were removed. To enable finer-scale functional comparisons, species-level GO slim summaries were subsequently aggregated across higher taxonomic levels. Graphical representations were created using ggplot2 (v4.0.0)^58^.

GO enrichment analyses were performed separately for each GO subontology — Biological Process (BP), Molecular Function (MF), and Cellular Component (CC) — using the enricher function from the clusterProfiler R package (v4.6.2)^59^. For each species, background gene sets were defined independently for each GO subontology as the set of all genes from that species with at least one GO full annotation in the corresponding subontology. Input gene sets consisted of retrocopied genes with at least one GO annotation in the corresponding subontology. Enrichment tests were conducted using species-specific background gene sets, and multiple testing correction was applied using the Benjamini-Hochberg procedure. Only GO terms with an adjusted p-value ≤ 0.05, gene set sizes between 10 and 500 genes, and fold enrichment ≥ 2 were retained for downstream analyses.

### Sex chromosome-autosome retrocopy trafficking analysis

To characterise retrocopy trafficking between sex chromosomes and autosomes, we classified all retrocopies according to the chromosomal location of their parental gene and their own genomic insertion site. Species were grouped by sex-determination system: XY (mammals, fishes, and reptiles) and ZW (birds and reptiles). For each species, parental genes and retrocopies were assigned to one of three compartments — X/Z chromosome, Y/W chromosome, or autosome — based on chromosome annotations provided by the VGP. All pairwise trafficking routes (e.g., X→Autosome, Autosome→X, X→Y) were quantified by counting the number of parental genes and retrocopies in each route. The retrocopy-per-parental ratio was computed for each route as the total number of retrocopies divided by the number of parental genes. To test whether specific trafficking routes showed significantly elevated or reduced retrocopy-per-parental ratios relative to the genomic background, Pearson’s chi-square tests of independence were performed comparing the parental-to-retrocopy ratio of each route against the corresponding ratio for all remaining routes combined. The relative size of each sex chromosome as a proportion of the total autosomal DNA content was estimated from genome assembly statistics to contextualise the magnitude of incoming retrocopy fluxes. Functional characterization of trafficking routes was performed by stratifying retrocopies according to conservation status (conserved vs. species-specific), coding potential (complete ORF retention), and selection signatures (purifying or positive selection based on dN/dS), and comparing the proportional representation of each category between sex-to-autosome and autosome-to-sex directions.

### Definition and classification of cancer-associated genes

Cancer-associated genes were defined based on the curated catalogue of Vogelstein et al. (2013)^36^, which identifies genes recurrently mutated in human cancers. Genes were classified as tumor suppressor genes (TSGs; n = 66) or oncogenes (n = 41) according to their established roles in tumorigenesis (**Supplementary Table 21**). For each VGP species, we identified retrocopies derived from cancer gene orthologs by mapping parental gene identifiers to orthology relationships obtained from the NCBI Gene database (https://ftp.ncbi.nlm.nih.gov/gene/DATA/gene_orthologs.gz) containing the corresponding human cancer genes. The proportion of cancer genes with at least one retrocopy was computed per lineage and compared to the genome-wide baseline-defined as the proportion of protein-coding genes with at least one retrocopy-using Fisher’s exact test. To quantify the relative accumulation of TSG versus oncogene retrocopies, we computed a TSG-bias index for each species, defined as (TSG − oncogene) / (TSG + oncogene), where TSG and oncogene refer to the total number of retrocopies normalized by the total number of retrocopied genes within each class. Index values above zero indicate preferential accumulation of TSG retrocopies; values below zero indicate oncogene bias. The number of mammalian orders in which each cancer gene harboured retrocopies was compared between TSGs and oncogenes using the two-sided Wilcoxon rank-sum test.

### Cancer mortality and longevity association analyses

Cancer mortality data were obtained from published zoo-based surveys^38,60^, which report the cumulative incidence of cancer as a cause of death across mammalian species maintained in zoological institutions. VGP species were cross-referenced with these datasets by taxonomic matching. Only mammalian orders with at least two species were further considered. For the longevity analysis, maximum lifespan data were compiled from the The Animal Ageing and Longevity (AnAge) database^61^. Within individual mammalian orders (Chiroptera, Primates, and Artiodactyla), the association between the TSG-bias index and maximum longevity was assessed using Spearman’s rank correlation. For the body mass analysis, VGP mammals were divided into three size categories (small, <10 kg; medium, 10–70 kg; large, >70 kg) based on published adult body mass estimates, and differences in TSG-bias index among categories were tested using the two-sided Wilcoxon rank-sum test.

### Comparisons with transposable elements

Transposable element (TE) annotations from RepeatModeler for each species were obtained from the NCBI repository via the UCSC Genome Browser VGP hub (https://hgdownload.soe.ucsc.edu/hubs/VGP/). To enable cross-species comparisons, we calculated the base pair coverage of each genome. The relative abundance of each TE family was expressed as the percentage of the genome it occupies, thereby normalizing TE content to genome size for each species.

### Phylogenetic generalized least squares

Phylogenetically corrected regressions were performed using the caper (v1.0.4)^62^ and ape (v5.8.1)^63^ R packages. A comparative data object was constructed with a variance-covariance matrix (vcv) derived from the phylogeny. Phylogenetic signal in the residuals was modelled using Pagel’s λ, which was estimated by maximum likelihood. Two models were fitted: (i) log_2_-transformed retrocopy abundance as a function of autonomous LINE content (L1/CR1/Tx1 elements), and (ii) log_2_-transformed retrocopy abundance as a function of SINE content. Model fit was assessed using adjusted R² and Akaike Information Criterion (AIC). To evaluate the contribution of phylogenetic structure, additional models were fitted with λ fixed at 1 (Brownian motion) and near zero (λ = 1×10⁻⁶, approximating ordinary least squares). Likelihood ratio tests comparing the maximum likelihood model to the λ ≈ 0 model were used to assess the significance of phylogenetic signal. All statistical analyses were conducted in R, and significance was evaluated using two-sided tests.

### RCPedia database and UCSC Genome Browser integration

All retrocopy and DNA-pseudogene annotations generated in this study were integrated into the RCPedia database^13^ (https://www.rcpedia.org/vgp). The database provides species-level annotations, including genomic coordinates, parental gene assignments, conservation status, ORF predictions, dN/dS-based selection classifications, and sex chromosome trafficking routes for all 244 VGP genomes^12^. Retrocopy annotations were additionally formatted as a custom track for the UCSC Genome Browser (genome.ucsc.edu), enabling visualisation of individual retrocopies in their genomic context alongside gene models, conservation scores, transposable element annotations, and regulatory data. Both resources are publicly accessible.

## Data availability

All the assemblies in the data sets presented here have been grouped under the VGP umbrella (NCBI BioProject PRJNA489243) and others: Darwin Tree of Life (DToL; PRJEB40665), European Reference Genome Atlas (ERGA; PRJEB43510), Bat1K (PRJNA489245), Cetacean Genomes Project 50 (PRJNA1020146), Denmark Yggdrasil (PRJNA955268), Genomics of Brazilian BioDiversity (GBB; PRJNA1180976), Oceanomics (PRJNA1046164), ATLASea (PRJEB64126), AfricaBP (PRJNA811786), and California Conservation Genomes Project (CCGP; PRJNA720569), among others. Raw data are also stored in GenomeArk (https://www.genomeark.org) along with intermediate files, annotations, and downstream analyses.

## Acknowledgements

We thank the Vertebrate Genomes Project (VGP) consortium for producing and sharing the genome assemblies and associated resources that formed the foundation of this study. We are also grateful to the many individuals and institutions who contributed to sample acquisition, access, curation, sequencing, genome annotation, and data generation, including field workers, permitting authorities, local collaborators, collection managers, sequencing centers, and technical teams. P.A.F.G. acknowledges support from the São Paulo Research Foundation (FAPESP; grants #2018/15579-8 and #2025/18246-3). M.J.O’C. thanks the Leverhulme Trust for support through a personal fellowship (RF-2024-492). R.L.V.M. acknowledges support from FAPESP (fellowship #2020/02413-4). D.M.M. acknowledges support from FAPESP (fellowship #2025/11174-7). R.L.V.M., G.D.A.G., N.D.R.D. and F.F.S. were supported by fellowships from the Young Scientist program, Hospital Sírio-Libanês. M.P.M.C acknowledges support from CNPq (142638/2023-4). G.A.S. acknowledges support from FAPESP (fellowship #23/11499-8). R.C.A.S. acknowledges support from FAPESP (scholarship #2025/12414-1). L.C.O. acknowledges support from FAPESP (scholarship #2024/13830-6).

## REFERENCES

1. Lynch, M. & Conery, J. S. The evolutionary demography of duplicate genes. J. Struct. Funct. Genomics 3, 35–44 (2003).

2. Kaessmann, H., Vinckenbosch, N. & Long, M. RNA-based gene duplication: mechanistic and evolutionary insights. Nat. Rev. Genet. 10, 19–31 (2009).

3. Navarro, F. C. P. & Galante, P. A. F. A Genome-Wide Landscape of Retrocopies in Primate Genomes. Genome Biol. Evol. 7, 2265–2275 (2015).

4. Carelli, F. N. et al. The life history of retrocopies illuminates the evolution of new mammalian genes. Genome Res. 26, 301–314 (2016).

5. Martinez-Morales, J. R. Toward understanding the evolution of vertebrate gene regulatory networks: comparative genomics and epigenomic approaches. Brief. Funct. Genomics 15, 315–321 (2016).

6. Dehal, P. & Boore, J. L. Two rounds of whole genome duplication in the ancestral vertebrate. PLoS Biol. 3, e314 (2005).

7. Schartl, M. et al. The genomes of all lungfish inform on genome expansion and tetrapod evolution. Nature 634, 96–103 (2024).

8. Böhne, A., Brunet, F., Galiana-Arnoux, D., Schultheis, C. & Volff, J.-N. Transposable elements as drivers of genomic and biological diversity in vertebrates. Chromosome Res. 16, 203–215 (2008).

9. Kikuta, H. et al. Genomic regulatory blocks encompass multiple neighboring genes and maintain conserved synteny in vertebrates. Genome Res. 17, 545–555 (2007).

10. Lynch, M. & Conery, J. S. The origins of genome complexity. Science 302, 1401–1404 (2003).

11. Kaessmann, H. Origins, evolution, and phenotypic impact of new genes. Genome Res. 20, 1313–1326 (2010).

12. Formenti, G., et al. The Vertebrate Genomes Project Phase I: A global reference genome resource. bioRxiv (2026) doi:10.64898/2026.06.24.732306.

13. Conceição, H. B. et al. RCPedia: a global resource for studying and exploring retrocopies in diverse species. Bioinformatics 40, (2024).

14. Wheeler, D. L. et al. Database resources of the National Center for Biotechnology Information. Nucleic Acids Res. 36, D13–21 (2008).

15. Song, Y. et al. A genomic compendium of hundreds of teleost fishes reveals their evolutionary landscape. Innovation (Camb.) 101177 (2025).

16. Uliano-Silva, M. et al. Elevated retrocopy burden and sloth-specific expansions illuminate mammalian genome evolution. BMC Biol. (2026) doi:10.1186/s12915-026-02632-5.

17. Beck, C. R. et al. LINE-1 retrotransposition activity in human genomes. Cell 141, 1159–1170 (2010).

18. Bruno, M. et al. Young KRAB-zinc finger gene clusters are highly dynamic incubators of ERV-driven genetic heterogeneity in mice. Nat. Commun. 16, 9608 (2025).

19. Derous, D., Sahu, J., Douglas, A., Lusseau, D. & Wenzel, M. Comparative genomics of cetartiodactyla: energy metabolism underpins the transition to an aquatic lifestyle. Conserv. Physiol. 9, coaa136 (2021).

20. Morales, A. E. et al. Bat genomes illuminate adaptations to viral tolerance and disease resistance. Nature 638, 449–458 (2025).

21. Huelsmann, M. et al. Genes lost during the transition from land to water in cetaceans highlight genomic changes associated with aquatic adaptations. Sci. Adv. 5, eaaw6671 (2019).

22. Nergadze, S. G., Rocchi, M., Azzalin, C. M., Mondello, C. & Giulotto, E. Insertion of telomeric repeats at intrachromosomal break sites during primate evolution. Genome Res. 14, 1704–1710 (2004).

23. Li, C. et al. Landscape of A-I RNA editing in mouse, pig, macaque, and human brains. Nucleic Acids Res. 53, gkaf534 (2025).

24. Turner, J. M. A. Meiotic sex chromosome inactivation. Development 134, 1823–1831 (2007).

25. Vibranovski, M. D., Lopes, H. F., Karr, T. L. & Long, M. Stage-specific expression profiling of Drosophila spermatogenesis suggests that meiotic sex chromosome inactivation drives genomic relocation of testis-expressed genes. PLoS Genet. 5, e1000731 (2009).

26. McLysaght, A. Evolutionary steps of sex chromosomes are reflected in retrogenes. Trends Genet. 24, 478–481 (2008).

27. Formenti, G., et al. The complete genome of a songbird. bioRxivorg (2025) doi:10.1101/2025.10.14.682431.

28. McCarrey, J. R. & Thomas, K. Human testis-specific PGK gene lacks introns and possesses characteristics of a processed gene. Nature 326, 501–505 (1987).

29. Potrzebowski, L. et al. Chromosomal gene movements reflect the recent origin and biology of therian sex chromosomes. PLoS Biol. 6, e80 (2008).

30. Uliano-Silva, M., et al. Retrocopy formation and domestication shape genome evolution in sloths and other xenarthrans. bioRxiv (2025) doi:10.1101/2025.09.25.678567.

31. Cason, A. L. et al. X-linked spermine synthase gene (SMS) defect: the first polyamine deficiency syndrome. Eur. J. Hum. Genet. 11, 937–944 (2003).

32. de Alencastro, G. et al. New SMS mutation leads to a striking reduction in spermine synthase protein function and a severe form of Snyder-Robinson X-linked recessive mental retardation syndrome. J. Med. Genet. 45, 539–543 (2008).

33. Pink, R. C. et al. Pseudogenes: pseudo-functional or key regulators in health and disease? RNA 17, 792–798 (2011).

34. Tollis, M., Schneider-Utaka, A. K. & Maley, C. C. The evolution of human cancer gene duplications across mammals. Mol. Biol. Evol. 37, 2875–2886 (2020).

35. Matthews, S., Nikoonejad Fard, V., Tollis, M. & Seoighe, C. Variable gene copy number in cancer-related pathways is associated with cancer prevalence across mammals. Mol. Biol. Evol. 42, msaf056 (2025).

36. Vogelstein, B. et al. Cancer genome landscapes. Science 339, 1546–1558 (2013).

37. Vazquez, J. M. & Lynch, V. J. Pervasive duplication of tumor suppressors in Afrotherians during the evolution of large bodies and reduced cancer risk. Elife 10, (2021).

38. Vincze, O. et al. Cancer risk across mammals. Nature 601, 263–267 (2022).

39. Sulak, M. et al. TP53 copy number expansion is associated with the evolution of increased body size and an enhanced DNA damage response in elephants. Elife 5, (2016).

40. Abegglen, L. M. et al. Potential Mechanisms for Cancer Resistance in Elephants and Comparative Cellular Response to DNA Damage in Humans. JAMA 314, 1850–1860 (2015).

41. Poliseno, L. et al. A coding-independent function of gene and pseudogene mRNAs regulates tumour biology. Nature 465, 1033–1038 (2010).

42. Vazquez, J., Kraft, M. & Lynch, V. A *CDKN2C* retroduplication in Bowhead whales is associated with the evolution of extremely long lifespans and alerted cell cycle dynamics. bioRxiv (2022) doi:10.1101/2022.09.07.506958.

43. Vincze, O. et al. Advancing cancer research via comparative oncology. Nat. Rev. Cancer 25, 740–748 (2025).

44. Lynch, V. J. Peto’s paradox revisited (revisited, revisited, revisited, and revisited yet again). Proc. Natl. Acad. Sci. U. S. A. 122, e2502696122 (2025).

45. Pertea, G. & Pertea, M. GFF Utilities: GffRead and GffCompare. F1000Research 9, 304 (2020).

46. Kiełbasa, S. M., Wan, R., Sato, K., Horton, P. & Frith, M. C. Adaptive seeds tame genomic sequence comparison. Genome Res. 21, 487–493 (2011).

47. Csárdi, G., et al. Igraph for R: R Interface of the Igraph Library for Graph Theory and Network Analysis. (Zenodo, 2026). doi:10.5281/ZENODO.7682609.

48. Haas, B. J. et al. De novo transcript sequence reconstruction from RNA-seq using the Trinity platform for reference generation and analysis. Nat. Protoc. 8, 1494–1512 (2013).

49. Thompson, J. D., Higgins, D. G. & Gibson, T. J. CLUSTAL W: improving the sensitivity of progressive multiple sequence alignment through sequence weighting, position-specific gap penalties and weight matrix choice. Nucleic Acids Res. 22, 4673–4680 (1994).

50. Suyama, M., Torrents, D. & Bork, P. PAL2NAL: robust conversion of protein sequence alignments into the corresponding codon alignments. Nucleic Acids Res. 34, W609–12 (2006).

51. Yang, Z. PAML 4: phylogenetic analysis by maximum likelihood. Mol. Biol. Evol. 24, 1586–1591 (2007).

52. Danis, C. et al. Inhibition of tau neuronal internalization using anti-tau single domain antibodies. Nat. Commun. 16, 3162 (2025).

53. Ho, L. si T. & Ané, C. A linear-time algorithm for Gaussian and non-Gaussian trait evolution models. Syst. Biol. 63, 397–408 (2014).

54. Carlson, M. org.Hs.eg.db. (Bioconductor, 2017). doi:10.18129/B9.BIOC.ORG.HS.EG.DB.

55. Gene Ontology Consortium. The Gene Ontology knowledgebase in 2026. Nucleic Acids Res. 54, D1779–D1792 (2026).

56. Greene, D., Richardson, S. & Turro, E. ontologyX: a suite of R packages for working with ontological data. Bioinformatics 33, 1104–1106 (2017).

57. Carlson, M. GO.db. (Bioconductor, 2017). doi:10.18129/B9.BIOC.GO.DB.

58. Wickham, H. ggplot2: Elegant Graphics for Data Analysis. (Springer International Publishing, 2016).

59. Wu, T. et al. clusterProfiler 4.0: A universal enrichment tool for interpreting omics data. Innovation (Camb.) 2, 100141 (2021).

60. Vincze, O. et al. Immunological surveillance against cancer across mammals. Nat. Commun. 16, 10333 (2025).

61. de Magalhães, J. P. et al. Human Ageing Genomic Resources: updates on key databases in ageing research. Nucleic Acids Res. 52, D900–D908 (2024).

62. Tyler, A., et al. Cape: Combined analysis of pleiotropy and epistasis for diversity outbred mice. CRAN: Contributed Packages The R Foundation 10.32614/cran.package.cape (2013).

63. Paradis, E. & Schliep, K. ape 5.0: an environment for modern phylogenetics and evolutionary analyses in R. Bioinformatics 35, 526–528 (2019).

