## Extended Figures for "Processed pseudogenes as dynamic substrates of vertebrate genome evolution"

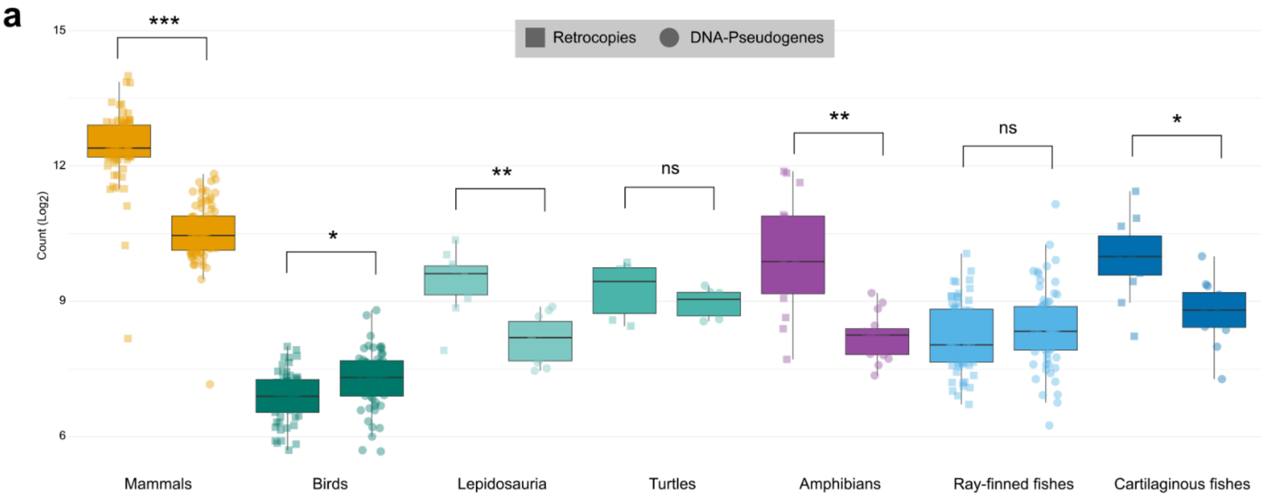


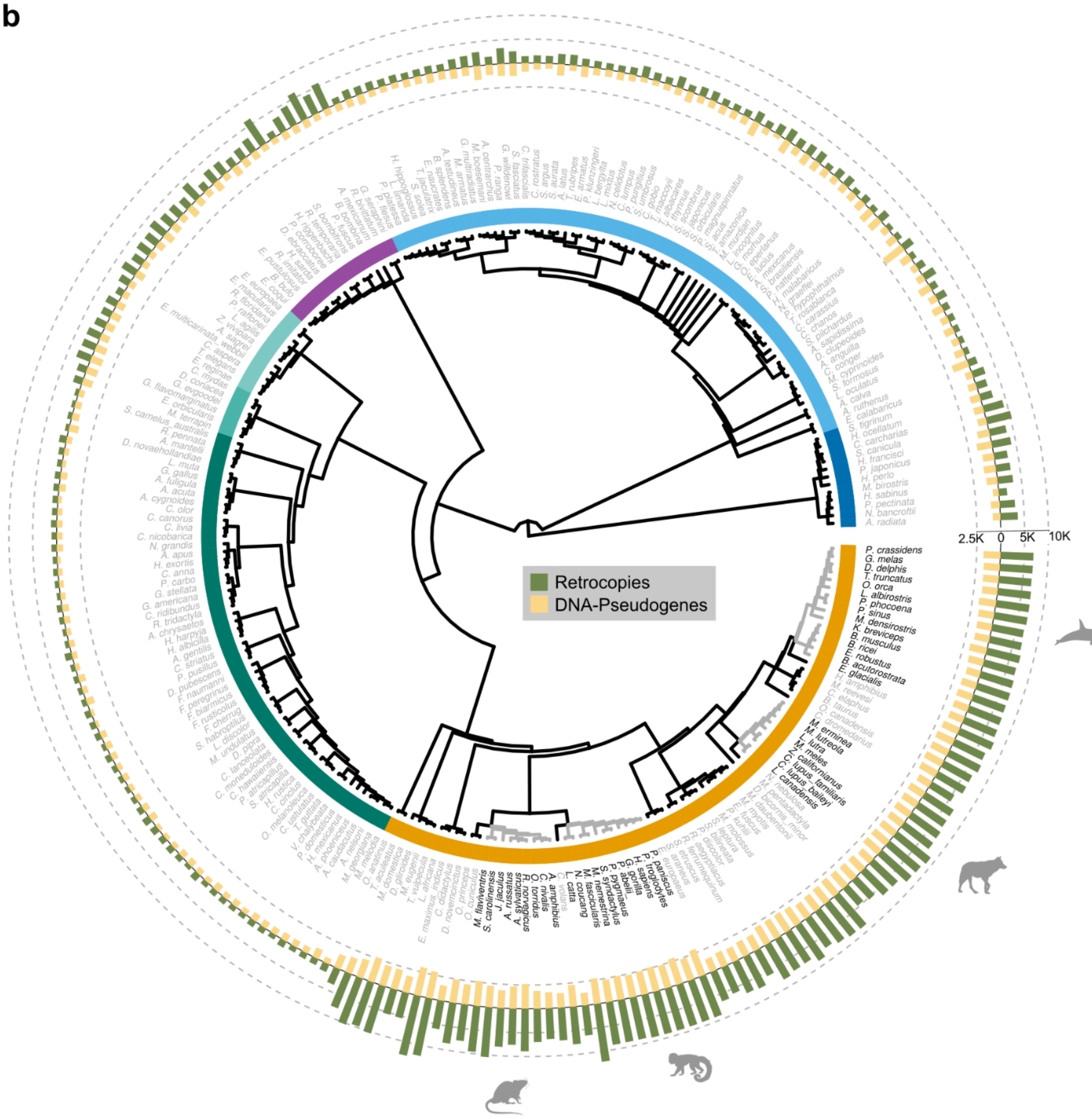


**Extended Fig 1. Genomic composition and phylogenetic distribution of retrocopies and pseudogenes across 244 vertebrate species. a,** Distribution of retrocopies and pseudogenes across vertebrate lineages. Box plots show

$log_{10}$-transformed counts per species within each taxonomic group, with retrocopies displayed as circles and pseudogenes as rectangles. Statistical significance determined by Wilcoxon rank-sum test: ***$P < 0.001$; **$P < 0.01$; *$P < 0.05$; ns, not significant. **b,** Phylogenetic distribution of retrocopies and pseudogenes across 244 vertebrate species. The cladogram represents phylogenetic relationships (branch lengths not scaled to evolutionary distance). The inner colored ring indicates taxonomic groups (mammals in orange, birds in purple, reptiles in turquoise/green, amphibians in pink, ray-finned fishes in light blue, cartilaginous fishes in dark blue). The outer stacked bar chart displays counts of pseudogenes (yellow) and retrocopies (green) for each species on a scale from 0 to 10,000. Species names are listed around the circumference.

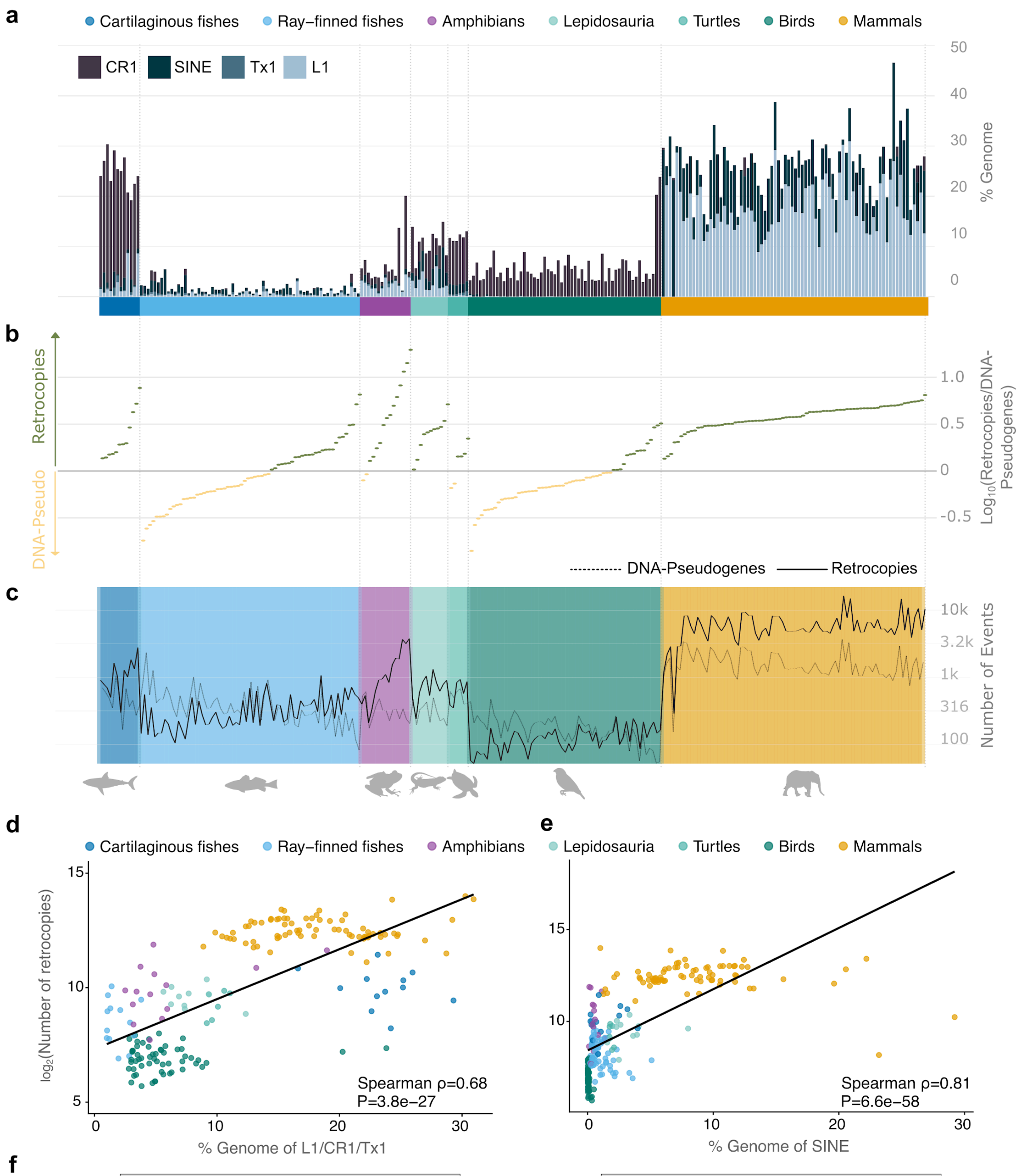


**f**

| PGLS model: Retrocopies ~ LINE (L1/CR1/Tx1) | | | |
|---|---|---|---|
| $\lambda$=0.966 | $\beta$=0.0359 | $R^2$=0.02331 | P=0.00969 |

| PGLS model: Retrocopies ~ SINE | | | |
|---|---|---|---|
| $\lambda$=0.975 | $\beta$=0.0362 | $R^2$=0.00723 | P=0.09735 |

**Extended Fig 2. Correlation between retrotransposon abundance and pseudogene formation across vertebrate evolution. a,** Genomic proportions (percentage of base pairs in the genome) for LINE/L1 elements (light blue), LINE/Tx1 elements (blue), SINE elements (darkest blue), and LINE/CR1 elements (dark brown). The colored bar beneath indicates taxonomic groups: cartilaginous fishes (dark blue), ray-finned fishes (light blue), amphibians (purple), reptiles (green/turquoise), and mammals (orange). **b,** Retrocopy-to-pseudogene ratio for each species across vertebrate genomes. Positive values indicate retrocopy (green) predominance and negative values indicate pseudogene (yellow) predominance. **c,** Corresponding counts of retrocopies (green line, left y-axis) and pseudogenes (yellow line, right y-axis) plotted on log-transformed scales. Each point represents a single species aligned with the retrotransposon data above. **d,** Correlation between the genomic fraction of autonomous non-LTR retrotransposons (L1, CR1, and Tx1 combined; % genome) and the number of retrocopies ($log_2$-transformed) across 244 vertebrate species (Spearman rho = 0.68, P = 3.8 × $10^{-27}$). **e,** Correlation between SINE content (% genome) and the number of retrocopies ($log_2$-transformed) across the same species (Spearman rho = 0.81, P = 6.6 × $10^{-58}$). In both panels, each point represents a single species, colored by taxonomic group; black lines indicate linear regression fits. **f,** Correlation between LINE/SINE content (% genome) and the number of retrocopies ($log_2$-transformed) across the species, accounting for phylogeny using Phylogenetic Generalised Least Squares (PGLS).

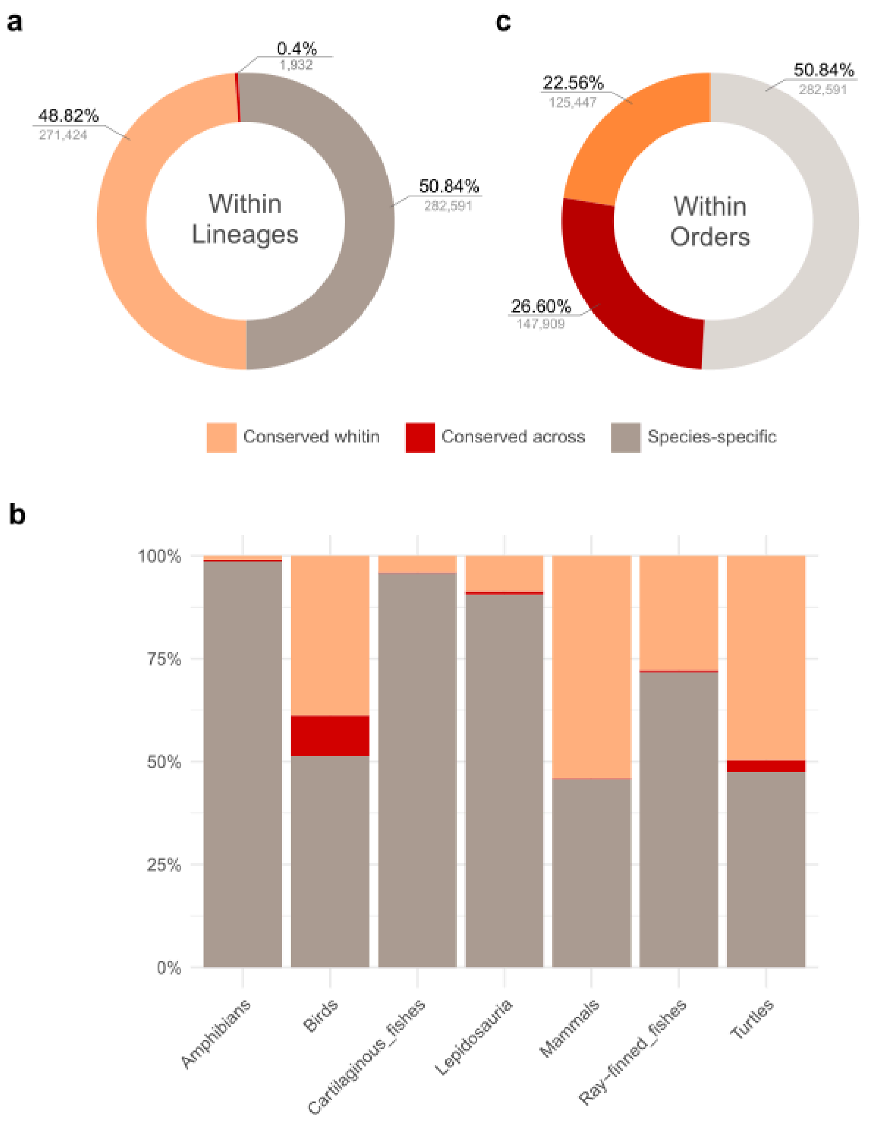


**Extended Fig 3. Conservation patterns of retrocopies across vertebrate lineages. a,** Overall distribution of retrocopies according to their conservation status: species-specific retrocopies, retrocopies conserved within the same lineage (shared among species belonging to the same lineage), and retrocopies conserved across different lineages (shared among species from distinct major lineages). **b,** Proportional distribution of these same conservation categories across each major vertebrate lineage, showing the relative contribution of species-specific retrocopies, retrocopies conserved within lineage, and retrocopies conserved across lineages. **c,** Overall distribution of retrocopies according to their conservation status: species-specific retrocopies, retrocopies conserved within the same order.

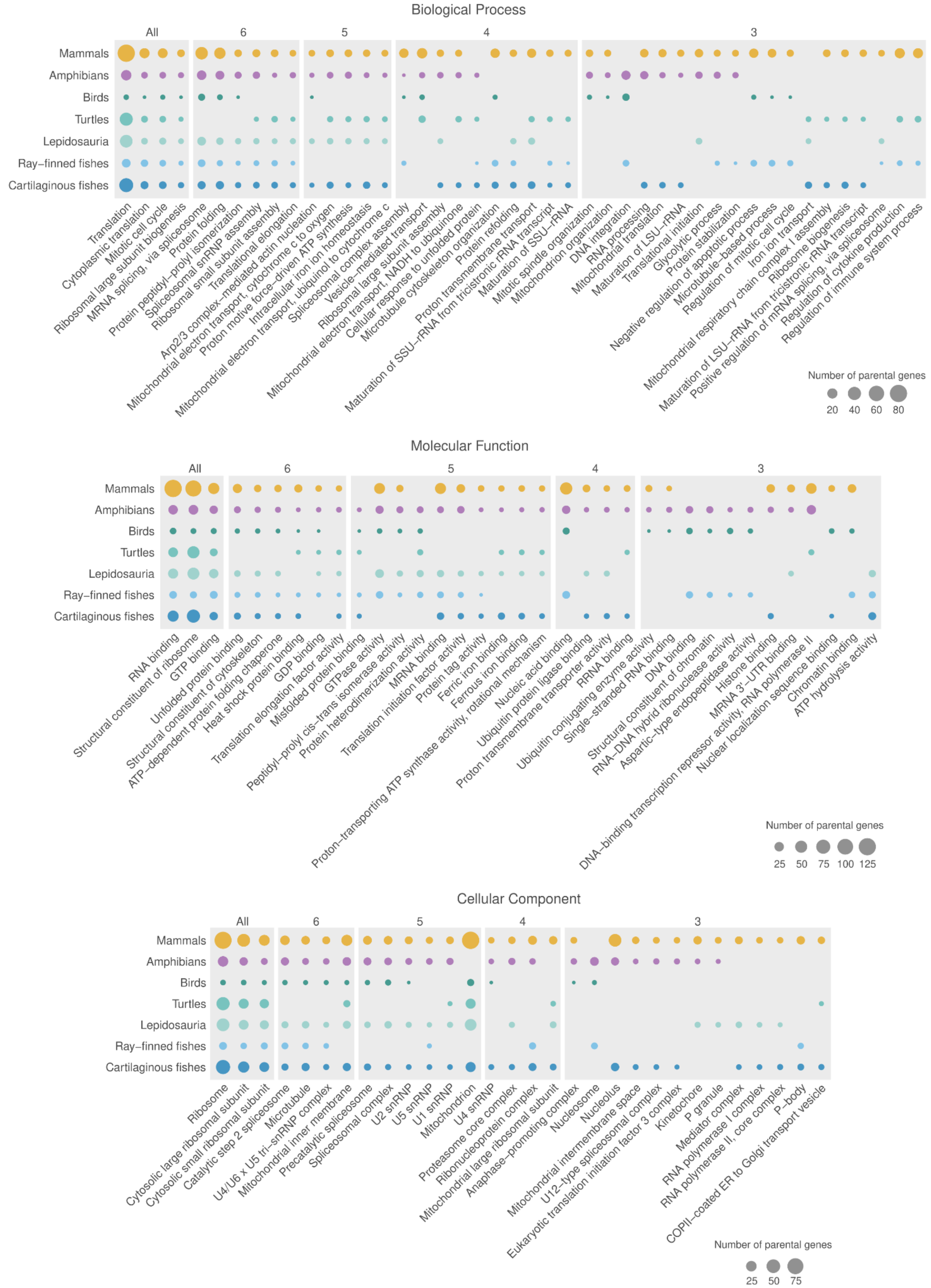


**Extended Fig 4. Hierarchical Gene Ontology enrichment of retrocopy parental genes, shared across vertebrate lineages.** GO terms significantly enriched (adjusted P-value < 0.05) in retrocopy parental genes, stratified by the number of vertebrate lineages sharing each enrichment. Panels display Biological Process (top), Molecular Function (middle), and Cellular Component (bottom). Column headers indicate the number of lineages (All = 7 lineages; 6, 5, 4, 3 = progressively fewer lineages).

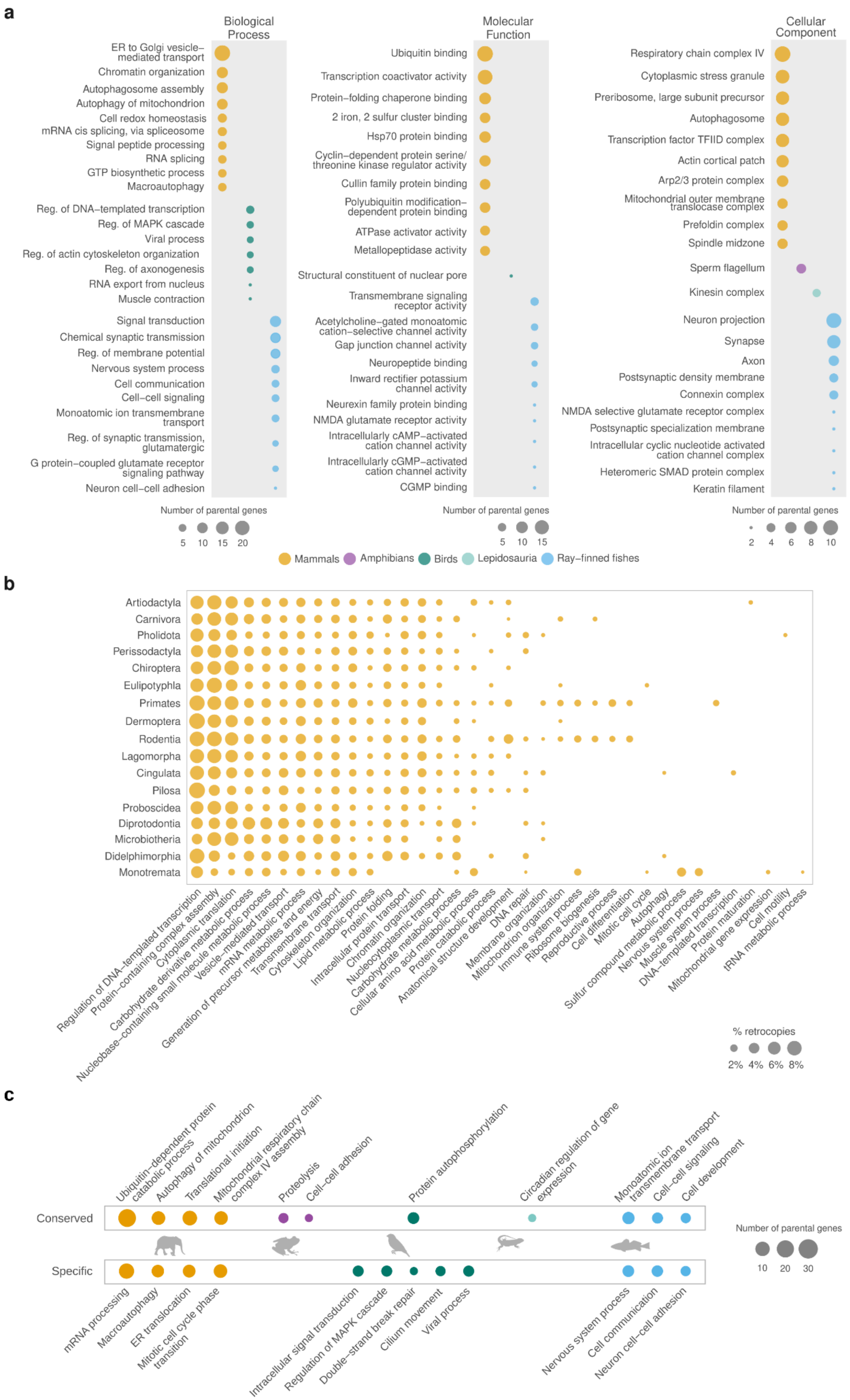


**Extended Fig. 5. Lineage-specific Gene Ontology enrichment of retrocopy parental genes. a,** GO terms significantly enriched (adjusted P-value < 0.05) exclusively within individual vertebrate lineages. Panels display Biological Process (top), Molecular Function (middle), and Cellular Component (bottom). Only lineages with significant lineage-restricted enrichments are shown for each GO category. **b,** Biological Process (GO slim) annotations across mammalian orders. Only GO terms associated with parental genes accounting for ≥ 1% of retrocopies in the order are shown. **c,** Gene Ontology (Biological Process) enrichment of parental genes giving rise to conserved (top) and species-specific (bottom) retrocopies (FDR < 0.05). Dot size reflects the number of parental genes; silhouettes indicate the lineages contributing to each enriched term.

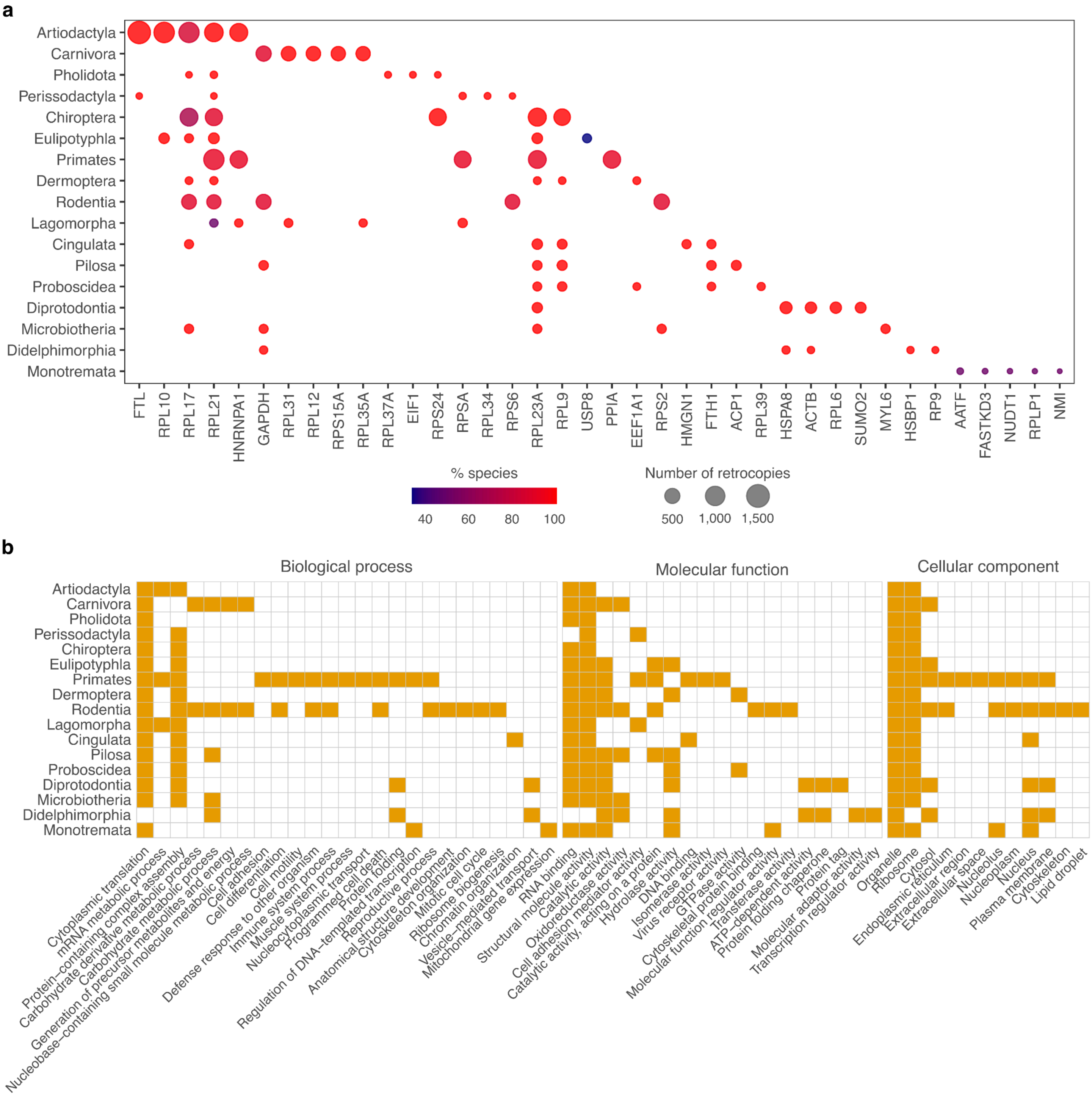


**Extended Figure 6. Number of retrocopies, frequency in the species, and functional annotation of the most frequently retrocopied genes across mammal orders. a,** Top retrocopied parental genes across mammalian orders, arranged phylogenetically. **b,** Gene Ontology slim annotations (Biological Process, Cellular Component, Molecular Function) of the top retrocopied parental genes in each mammalian order.

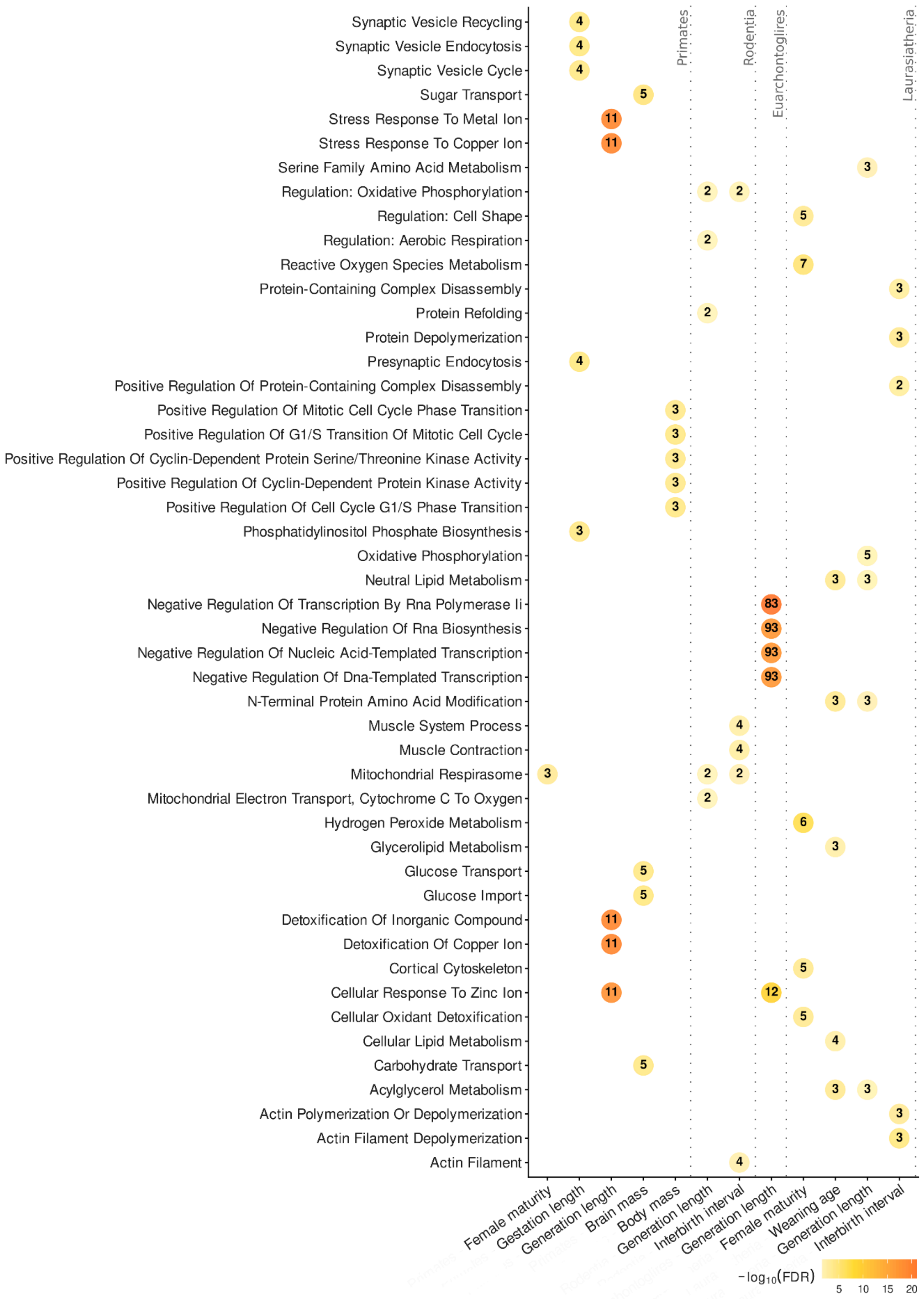


**Extended Fig 7. Retrocopy fraction and life-history trait associations reveal order-specific functional signatures in mammals.** Gene Ontology (Biological Process) enrichment of orthogroups associated with each trait. Only significantly enriched terms (FDR < 0.05) are shown.

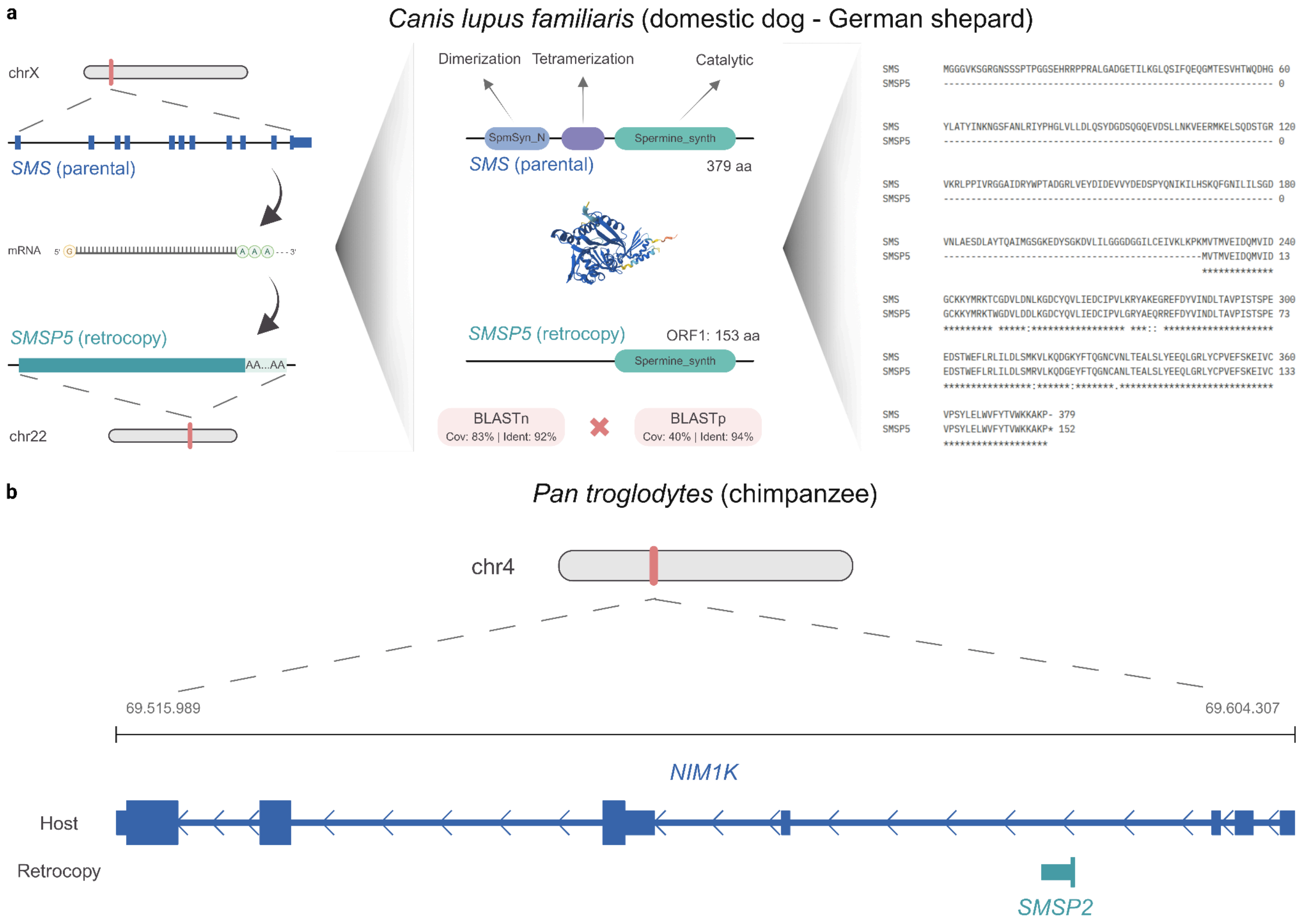


**Extended Fig 8. *SMS* retrocopies in domestic dog and chimpanzee. a,** *SMSP5*, located on chr22, is a retrocopy of the parental gene *SMS* (left panel). Alignment showing high identity in the C-terminal portion of the protein, according to BLAST (v2.17.0). ORF1 of the retrocopy conserves the catalytic spermine/spermidine synthase domain (right panel), according to HMMER (v3.4); **b,** *SMSP2* is another retrocopy of the parental gene *SMS*. This case is an intragenic (*NIM1K*) RTC located on chr4 in the chimpanzee. *SMS:* Spermine Synthase; *NIM1K*: *NIM1* serine/threonine protein kinase.

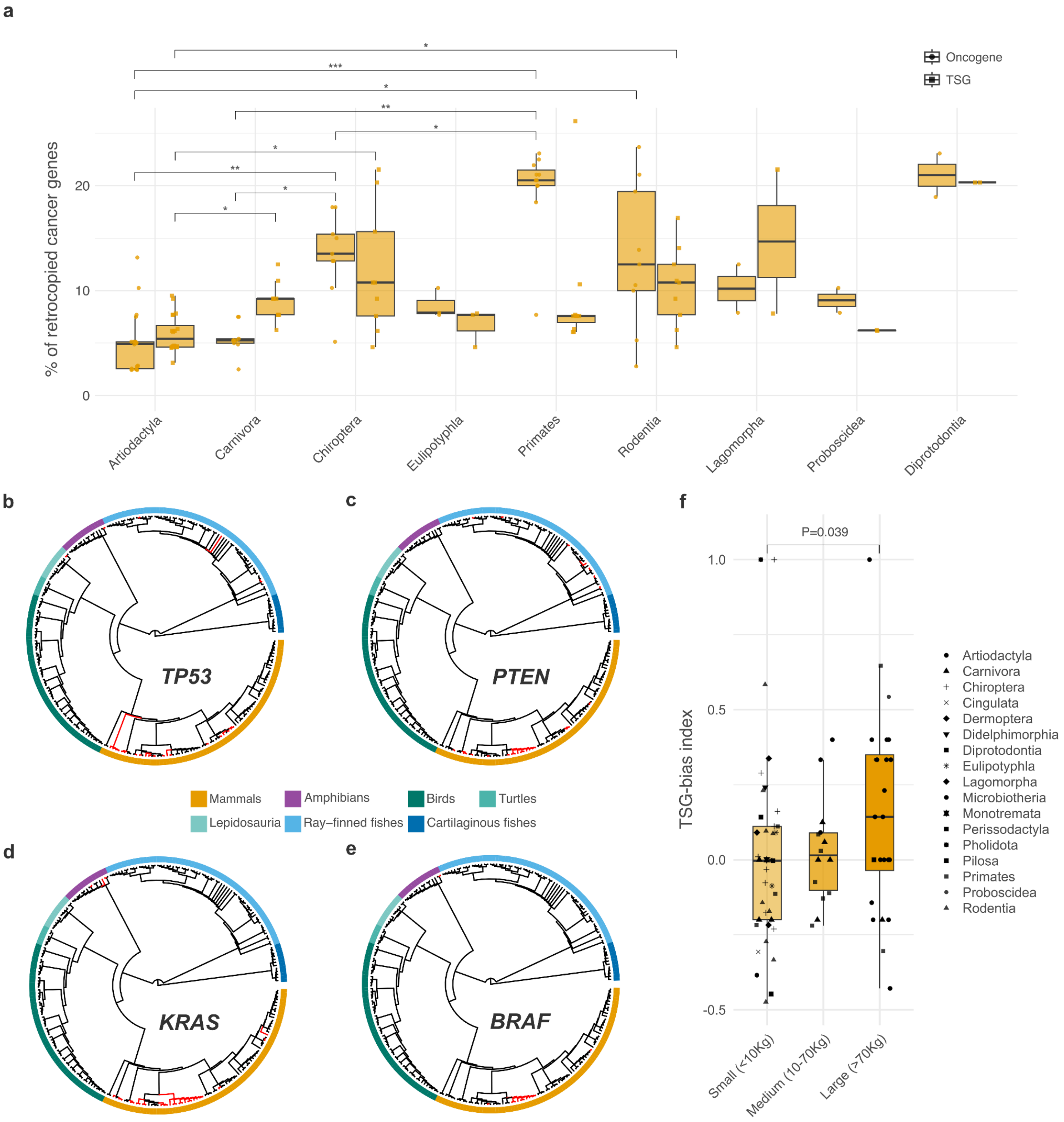


**Extended Fig 9. Phylogenetic distribution of cancer gene retrocopies and tumor suppressor bias across vertebrate lineages. a,** Proportion of tumor suppressor genes and oncogenes that have generated at least one retrocopy in at least one mammalian VGP species, stratified by order. Statistical significance determined by Wilcoxon rank-sum tests: * $p < 0.05$, ** $p < 0.01$, *** $p < 0.001$. **b-e**, Phylogenetic distribution of retrocopies derived from the cancer genes (b) *TP53*, (c) *PTEN*, (d) *KRAS*, and (e) *BRAF* across 244 vertebrate species. Cladograms represent phylogenetic relationships among species (branch lengths not scaled to evolutionary distance). Branches highlighted in red indicate species harboring at least one retrocopy derived from the corresponding parental gene. The inner colored ring indicates taxonomic groups (mammals in orange, amphibians in purple, birds in dark green, turtles in dark turquoise/green, lepidosauria in light turquoise/green, ray-finned fishes in light blue, cartilaginous fishes in dark blue). **f,** Distribution of TSG-bias indices across mammalian species stratified by body size categories (small, medium, and large). Values above zero indicate preferential accumulation of TSG retrocopies; values below zero indicate oncogene bias. Statistical significance determined by Wilcoxon rank-sum test, P=0.039.
