## Supplementary Material for "Processed pseudogenes as dynamic substrates of vertebrate genome evolution"

#### INDEX

#### Supplementary Tables

**Table captions**
**Supplemental Table 1.** List of analyzed clades, lineages, orders and species.
**Supplemental Table 2.** Number of retrocopies, pseudogenes, coding genes and transposable element classes (L1, CR1, Tx1 and SINEs) by species.
**Supplemental Table 3.** Number of conserved and species-specific retrocopies by species.
**Supplemental Table 4.** Conserved retrocopies across taxonomic lineages and orders, and comparison with protein-coding gene conservation (Fisher's exact tests).
**Supplemental Table 5.** Gene Ontology enrichment of retrocopy parental genes across the seven major vertebrate lineages (FDR < 0.05).

**Supplemental Table 6.** Gene Ontology Slim (Biological Process) annotations of retrocopy parental genes across the 17 mammals orders.
**Supplemental Table 7.** Gene Ontology (Biological Process) enrichment of parental genes giving rise to conserved and species-specific retrocopies (FDR < 0.05).
**Supplemental Table 8.** Most frequently retrocopied parental genes across vertebrate lineages.
**Supplemental Table 9.** Most frequently retrocopied parental genes across mammal's order.
**Supplemental Table 10.** Gene Ontology Slim annotations of the top retrocopied parental genes by mammals order.
**Supplemental Table 11.** Gene Ontology Slim (Biological Process) annotations of the top retrocopied parental genes by lineage.
**Supplemental Table 12.** Retrocopies with retained open reading frames evolving under selection by species.
**Supplemental Table 13.** Gene Ontology (Biological Process) annotations of parental genes giving rise to order-restricted retrocopies under selection in mammals.
**Supplemental Table 14.** Orthogroups with significant associations between retrocopy fraction and life-history traits across mammalian subgroups (log10-transformed, $q < 0.05$, occupancy ≥ 0.7). Empty lambda-model cells indicate that the Pagel's-λ model fit failed; the BM model fit successfully and was used for inference (best_model = BM).
**Supplemental Table 15.** Retrocopy movement between sex chromosomes (X, Y) and autosomes (A) by species.
**Supplemental Table 16.** Retrocopy movement between sex chromosomes (Z, W) and autosomes (A) by species.
**Supplemental Table 17.** Retrocopies per parental gene from chrX/Y, chrZ/W and all chromosomes.
**Supplemental Table 18.** *PGK1* retrocopies by mammals species.
**Supplemental Table 19.** *PDHA1* retrocopies by mammals species.
**Supplemental Table 20.** *SMS* retrocopies by mammals species.
**Supplemental Table 21.** List of cancer-associated genes.
**Supplemental Table 22.** Propensity of cancer genes for retrotransposition (Fisher's exact tests) across lineages.
**Supplemental Table 23.** Composition of retrocopied cancer genes by species.
**Supplemental Table 24.** Retrocopies of cancer genes by species.
**Supplemental Table 25.** TSG-bias index and longevity across mammalian species.

### Supplementary notes

#### Supplementary Note 1. Decoupling between retrotransposon abundance and retrocopy formation across vertebrates

A central observation of our analysis is that the strong cross-species association between non-LTR retrotransposon content and retrocopy abundance largely dissolves once phylogenetic non-independence is accounted for. Across the 244 vertebrate species analyzed, the genomic fraction occupied by autonomous LINEs (L1, CR1, and Tx1 combined) correlates strongly with retrocopy count under a phylogeny-naive Spearman test ($\rho = 0.68$, $P = 3.8 \times 10^{-27}$), and the association is even stronger for SINEs ($\rho = 0.81$, $P = 6.6 \times 10^{-58}$; **Extended Fig. 2d,e**). These coefficients reflect a broad mammalian outlier signal — mammals carry both the highest L1 content and the largest retrocopy reservoirs — and are consistent with the biochemical understanding that mature L1 ORF1/ORF2 proteins are the obligate enzymatic machinery for processed pseudogene formation in vertebrates[1]. Under Phylogenetic Generalized Least Squares (PGLS), however, the regression slopes shrink and the explained variance collapses to $R^2 \approx 0.023$ for LINEs ($\lambda = 0.966$, $\beta = 0.0359$, $P = 0.010$) and $R^2 = 0.007$ for SINEs ($\lambda = 0.975$, $\beta = 0.0362$, $P = 0.097$; **Extended Fig. 2f**). The high values of Pagel's $\lambda$ indicate that nearly all of the trait covariation tracks the phylogeny rather than direct, species-level co-variation between TE load and retrocopy yield.

We interpret this decoupling not as evidence against a mechanistic role for LINEs in retroduplication, but as quantitative support for the view that the rate-limiting steps of retrocopy formation lie downstream of retrotransposon DNA abundance. Bulk genomic content reflects the integrated history of TE insertion, expansion and deletion over hundreds of millions of years[2,3]; current retrocopy yield depends instead on (i) the fraction of L1 copies that remain transcriptionally and translationally competent in the germline[1], (ii) the availability of host cofactors and chaperones that license target-primed reverse transcription, and (iii) the germline expression profile of parental genes, which dictates substrate availability. These determinants vary across lineages largely independently of total TE load.

The same dissociation has been described in narrower mammalian comparative frameworks[5], and our pan-vertebrate sampling confirms it as a general feature of the group rather than a mammal-restricted pattern. The lineage-specific architecture of this dissociation is further illustrated by extreme outliers within mammals. The two monotremes in our dataset — *Ornithorhynchus anatinus* (289 retrocopies) and *Tachyglossus aculeatus* (1,207) — carry retrocopy loads one to two orders of magnitude below those of placentals and marsupials of comparable genome size and L1 content (**Supplementary Table 2; Supplementary Fig. 2**). This is consistent with reported reductions in active L1 sub-lineages in monotremes[2] and reinforces the conclusion that competent retrotransposition machinery, rather than retroelement abundance per se, governs retrocopy production.

Together, these observations motivate the two-stage model presented in the main text, in which a broadly conserved formation bias towards housekeeping parental genes is layered onto lineage-specific filters that determine which retrocopies are produced and which are retained in the genome.

#### Supplementary Note 2. Cancer gene retrocopies and Peto's paradox: interpretation and caveats

Our finding that tumor suppressor genes (TSGs) and oncogenes are retrocopied in 81.8% and 80.5% of the queried gene set, respectively (**Fig. 5a; Supplementary Table 22**), establishes cancer-relevant loci as a near-universal substrate of retroduplication across vertebrates. The biologically more interesting signal lies in the asymmetric distribution of these retrocopies — the TSG-bias index towards positive values in large-bodied, long-lived mammalian orders (Proboscidea, Artiodactyla,

Chiroptera) and the convergent amplification of *TP53* in elephants, together with the broad retrotransposition of *PTEN*, *KRAS* and *BRAF* across multiple orders (**Fig. 5b-e**; **Extended Fig. 9c-f**). Several caveats merit explicit discussion before this pattern can be invoked as a contributor to Peto's paradox[3,4].

First, the within-order longevity correlations are positive but statistically underpowered. In Chiroptera, the TSG-bias × maximum-lifespan Spearman coefficient is $\rho = 0.533$ with $P = 0.148$ ($n = 9$), and in Primates $\rho = 0.176$, $P = 0.632$ ($n = 10$). Neither remains significant after multiple-testing correction. We retain these analyses in the main text because the direction of effect is consistent across two phylogenetically distant orders with independent retrotransposition histories, but the inference at this stage is suggestive rather than confirmatory. Expansion of VGP sampling across Chiroptera and Primate genomes will be required to resolve whether the trend stabilizes or attenuates.

Second, the reference set is asymmetric: 66 TSGs versus 41 oncogenes, drawn from human cancer gene annotations[9]. A naive count ratio could therefore inflate apparent TSG bias. We verified that the TSG-bias index — defined as (TSG − oncogene)/(TSG + oncogene) retrocopies — remains qualitatively unchanged when normalized by reference-set size and when restricted to bidirectional ortholog-resolved gene sets (**Supplementary Table 23**). Nevertheless, the use of human-centric cancer gene classifications in a pan-vertebrate analysis carries unavoidable ascertainment bias, and we have framed all conclusions accordingly.

Third, the presence of a retrocopy does not entail function. The majority of cancer-gene-derived retrocopies across vertebrates lack intact open reading frames and evolve under neutrality (**Supplementary Fig. 9**); only a small minority carry the joint hallmarks of ORF retention and detectable purifying selection. Any functional contribution to anti-tumor phenotypes — through dosage compensation, ceRNA activity, or recruitment as protein-coding paralogs — must be demonstrated locus by locus. The *SMS* retrocopies described in domestic dog (*SMSP5*, chr22) and chimpanzee (*SMSP2*, intragenic to *NIM1K* on chr4; **Extended Fig. 8**) illustrate the kind of evidence required: ORF retention, conservation of the catalytic spermine/spermidine synthase domain detectable by HMMER profile search, and a genomic context compatible with expression. Generalizing from such individual cases to a lineage-level anti-cancer phenotype demands transcriptomic and functional follow-up beyond the scope of this resource paper.

Within these limits, our results are consistent with, and complementary to, the *TP53* expansion in Proboscidea[10,11], the *LIF6* recruitment in elephants[12], and the broader pattern of cancer-resistance gene duplication documented in Cetacea[13] and long-lived bats[14]. Retrotransposition emerges as an under-appreciated route for cancer-relevant copy-number variation, distinct from segmental duplication, whose contribution to organismal cancer susceptibility now warrants systematic functional dissection.

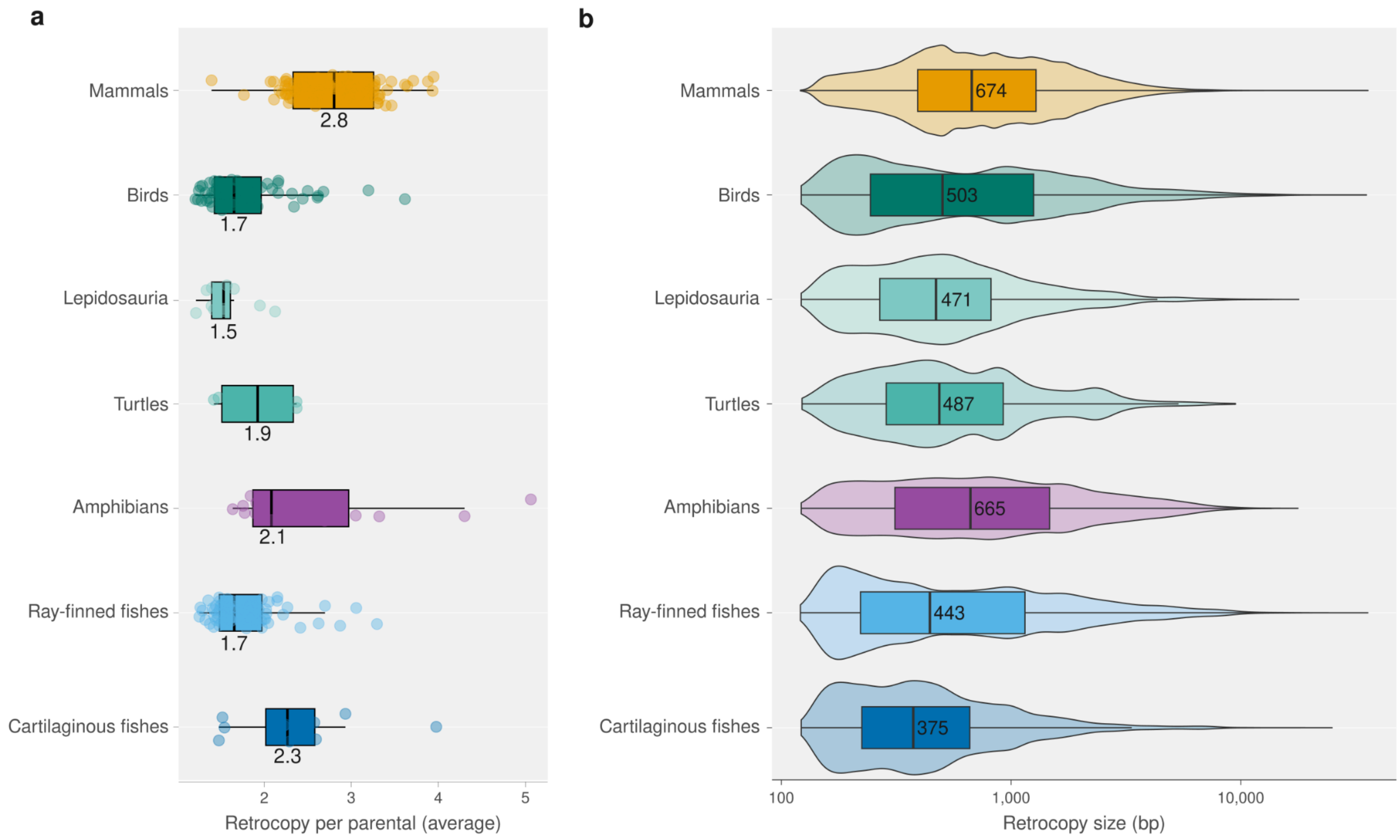

**Supplementary Fig 1. General features of retrocopies in vertebrates.**

**a**, Number of retrocopies per parental gene across the orders studied; each dot represents a species. **b**, Distribution of retrocopy lengths (bp) across the orders studied. Numbers indicate the median value for each plot.

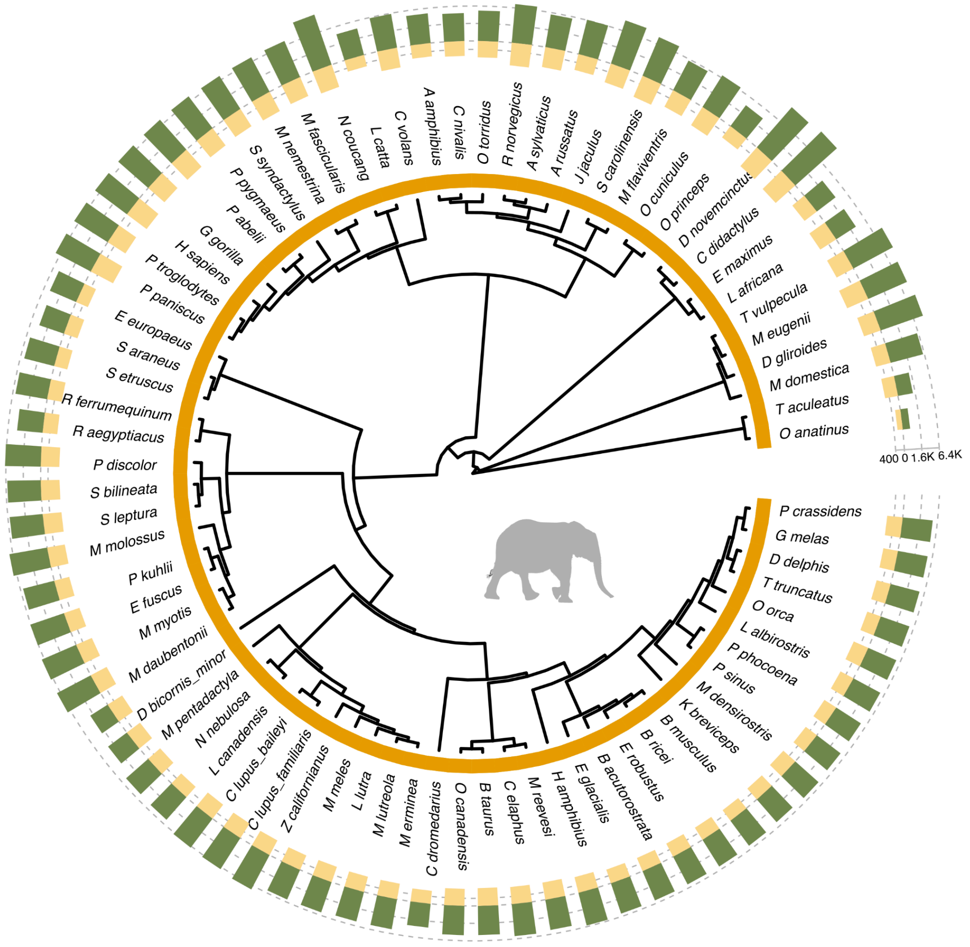

**Supplementary Fig 2. Phylogenetic distribution of retrocopies and pseudogenes across mammal species.**

The cladogram represents phylogenetic relationships with branch lengths not scaled to evolutionary distance. The outer stacked bar chart displays counts of retrocopies (green) and non-processed pseudogenes (yellow) for each species on a scale from 0 to 10,000. Species names are listed around the circumference.

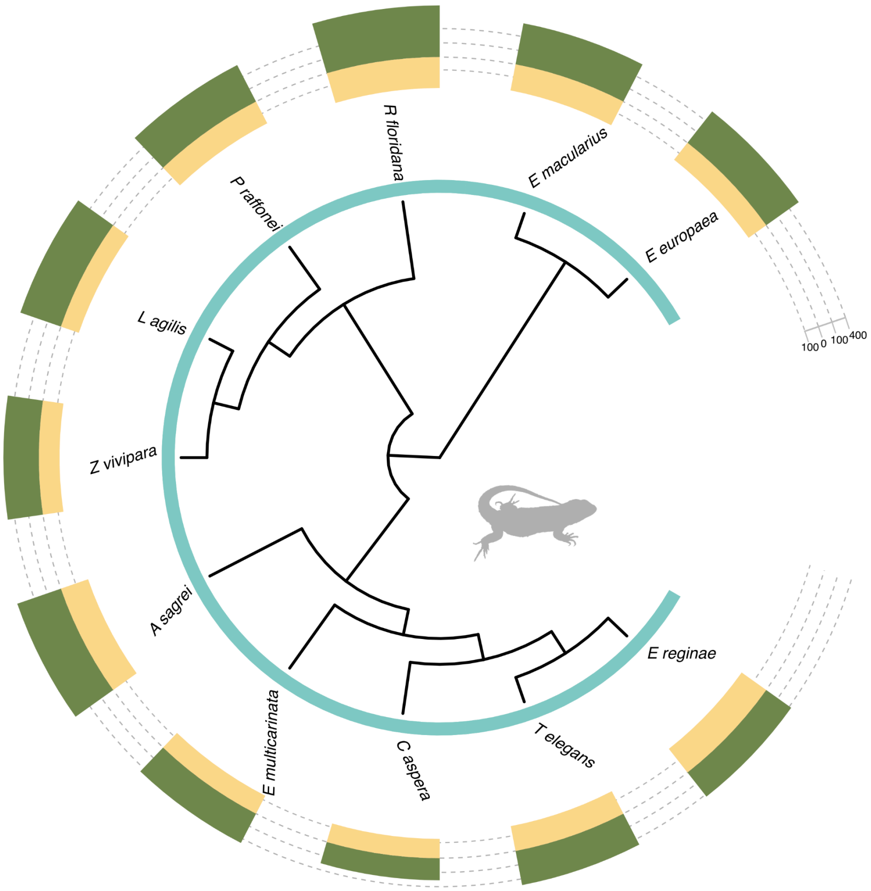

**Supplementary Fig 3. Phylogenetic distribution of retrocopies and pseudogenes across lepidosaurs species.** The cladogram represents phylogenetic relationships with branch lengths not scaled to evolutionary distance. The outer stacked bar chart displays counts of retrocopies (green) and non-processed pseudogenes (yellow) for each species on a scale from 0 to 10,000. Species names are listed around the circumference.

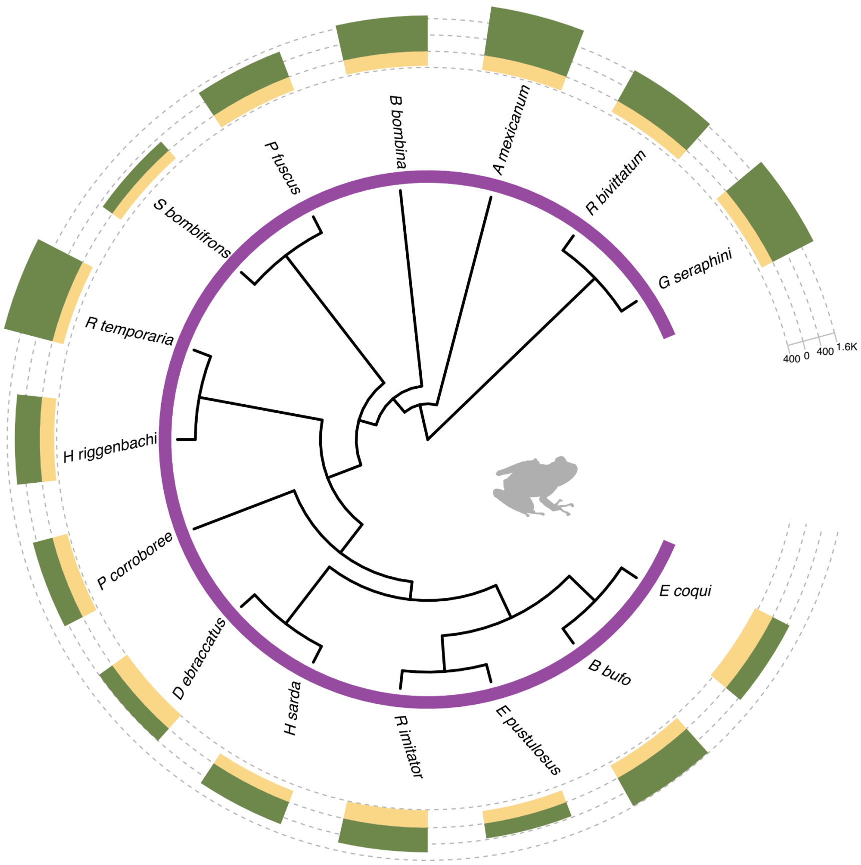

**Supplementary Fig 4. Phylogenetic distribution of retrocopies and pseudogenes across amphibian species.** The cladogram represents phylogenetic relationships with branch lengths not scaled to evolutionary distance. The outer stacked bar chart displays counts of retrocopies (green) and non-processed pseudogenes (yellow) for each species on a scale from 0 to 10,000. Species names are listed around the circumference.

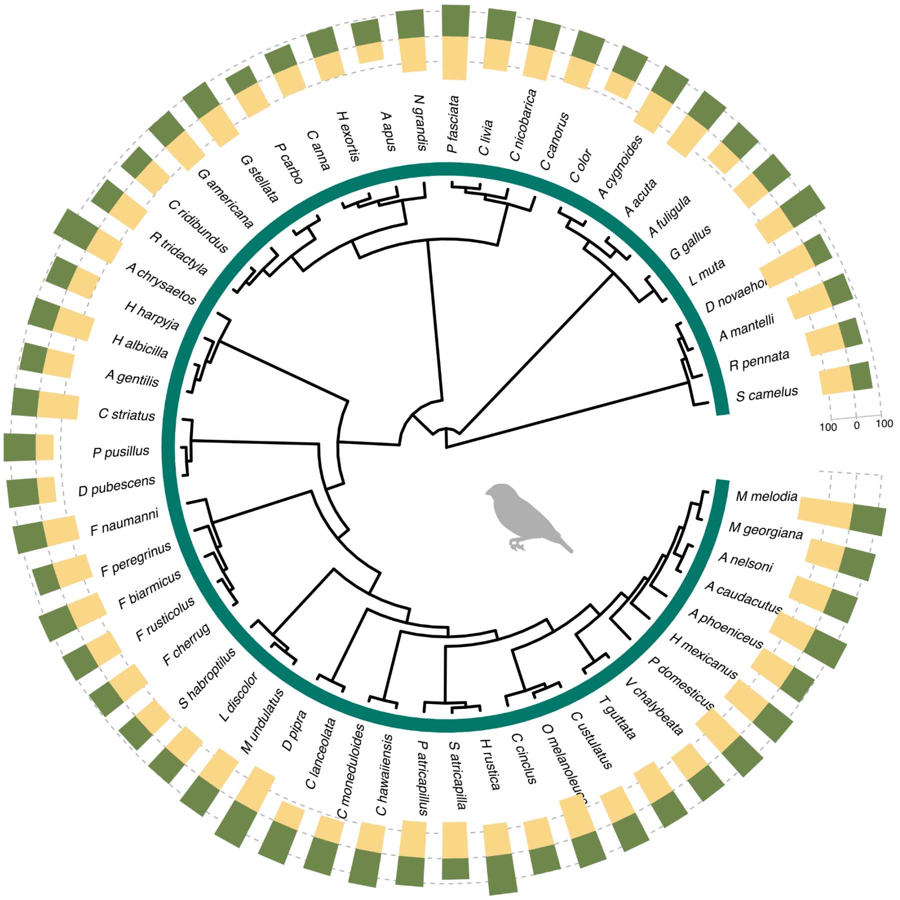

**Supplementary Fig 5. Phylogenetic distribution of retrocopies and pseudogenes across bird species.**

The cladogram represents phylogenetic relationships with branch lengths not scaled to evolutionary distance. The outer stacked bar chart displays counts of retrocopies (green) and non-processed pseudogenes (yellow) for each species on a scale from 0 to 10,000. Species names are listed around the circumference.

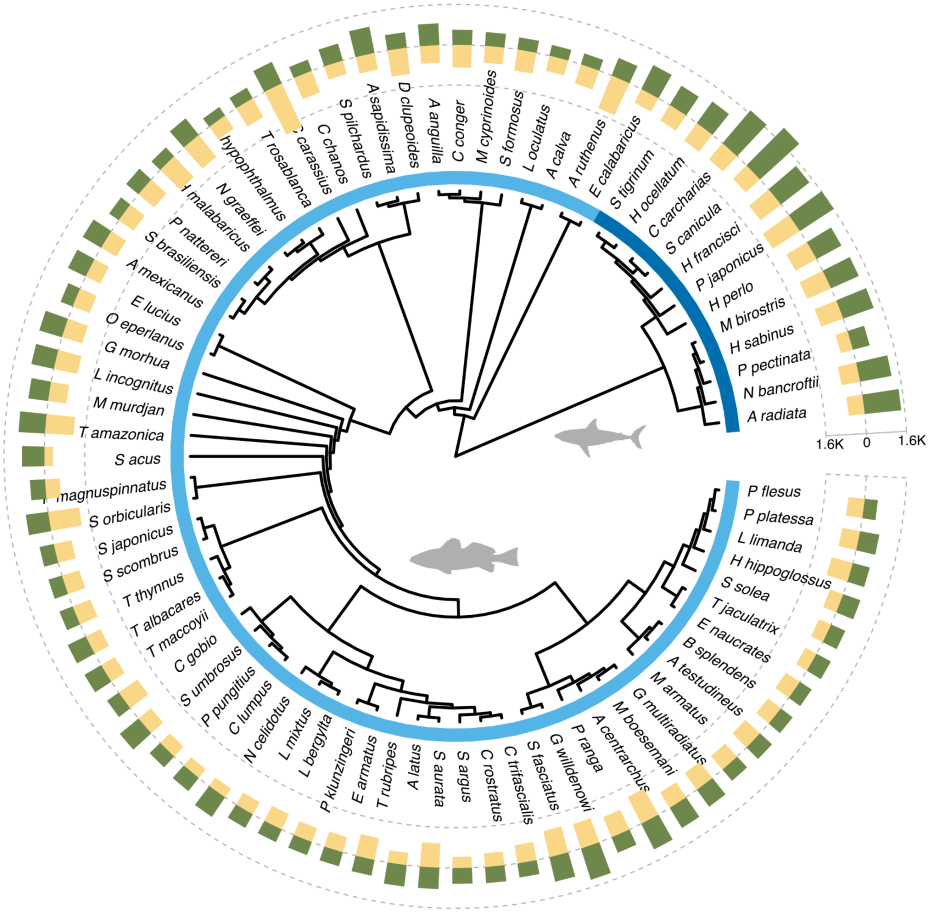

**Supplementary Fig 6. Phylogenetic distribution of retrocopies and pseudogenes across fish (ray-finned and cartilaginous fishes) species.**

The cladogram represents phylogenetic relationships with branch lengths not scaled to evolutionary distance. The outer stacked bar chart displays counts of retrocopies (green) and non-processed pseudogenes (yellow) for each species on a scale from 0 to 10,000. Species names are listed around the circumference. The inner colored ring indicates taxonomic groups (ray-finned fishes in light blue and cartilaginous fishes in dark blue).

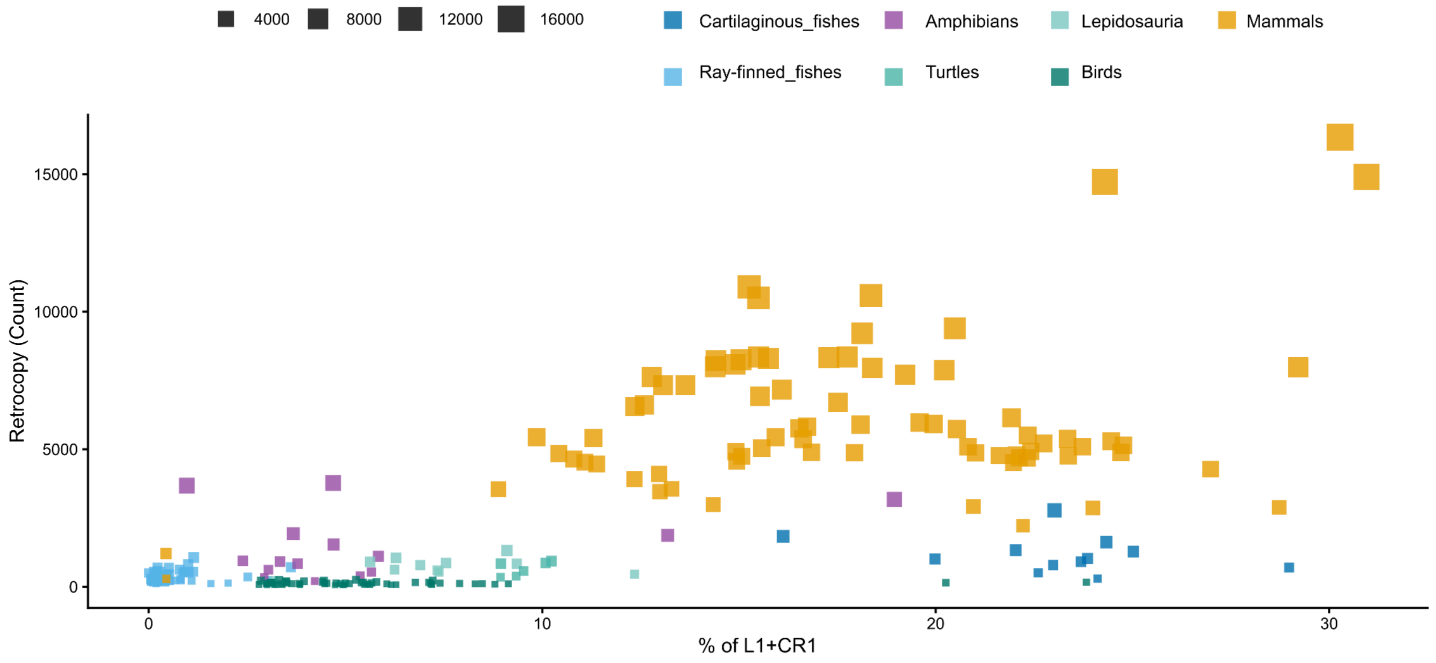

**Supplementary Fig 7. Percentage of L1 (LINE) plus CR1 (LINE) in the genome (bp) versus retrocopy count.**

Each square represents a species, color-coded by lineage, and the size refers to the number of retrocopies in that species genome.

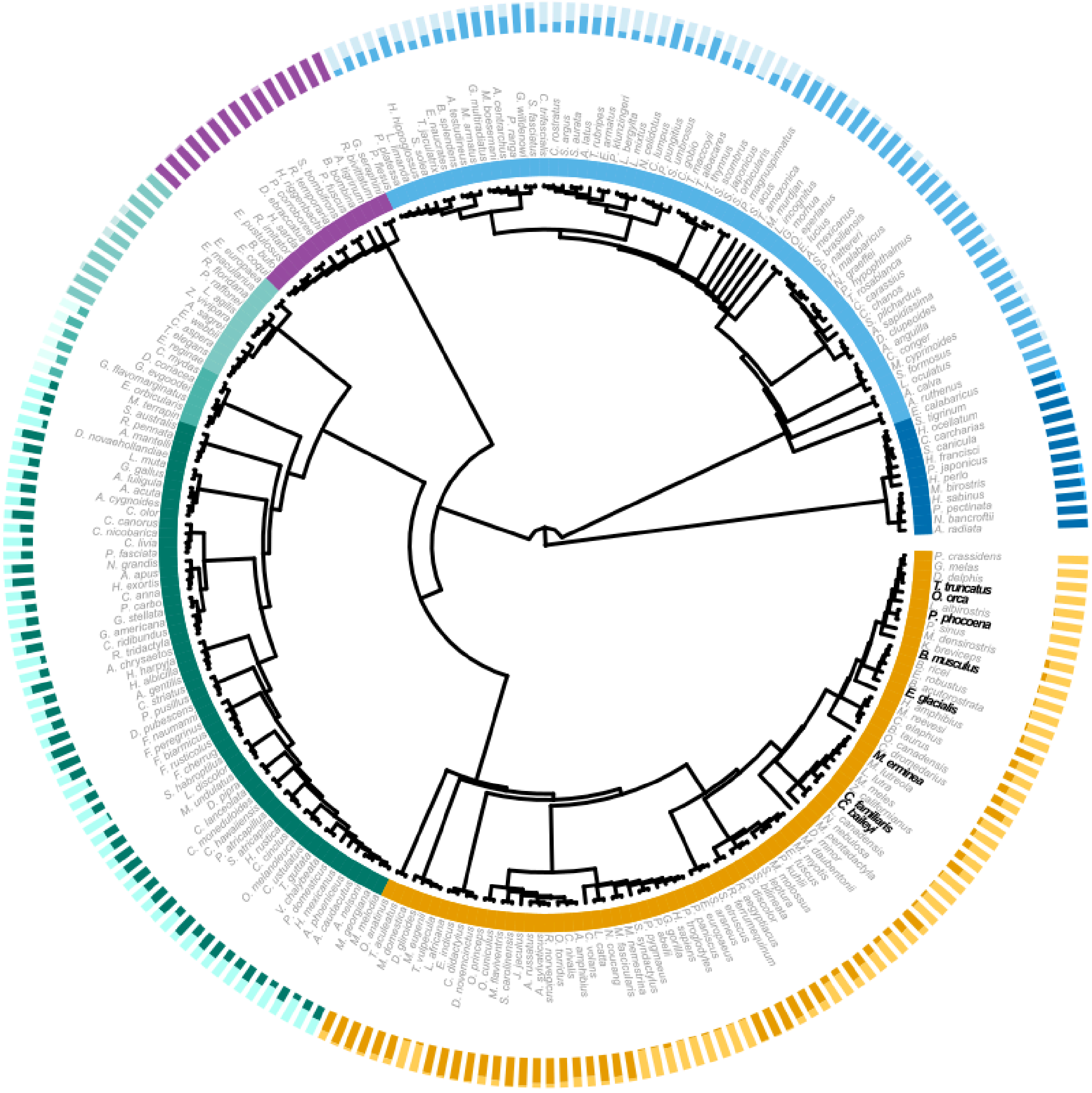

**Supplementary Fig 8. Species-level distribution of species-specific (dark) and conserved (light) retrocopies mapped onto the vertebrate phylogeny.**

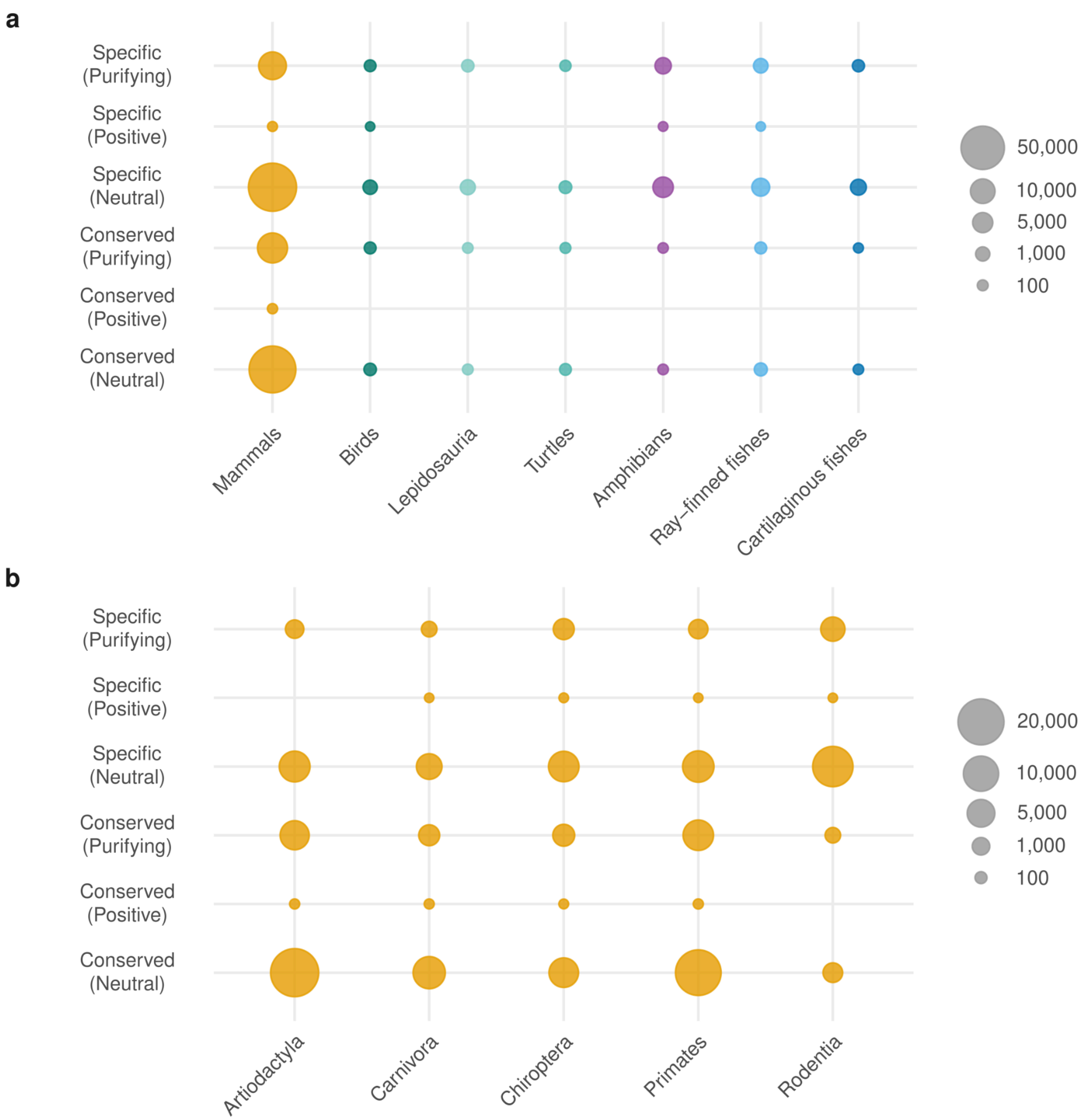

**Supplementary Fig 9. ORF-bearing retrocopies evolving under purifying, positive or neutral selection.**

**a**, Numbers of ORF-bearing retrocopies evolving under purifying, positive or neutral selection, stratified by conservation status (conserved vs. species-specific). Dot size reflects retrocopy count. **b,** As in **a,** resolved across major mammalian orders.

REFERENCES

1. Goodier, J. L. Retrotransposition in tumors and brains. *Mob. DNA* **5**, 11 (2014).
2. Zhou, Y. *et al.* Platypus and echidna genomes reveal mammalian biology and evolution. *Nature* **592**, 756–762 (2021).
3. Lynch, V. J. Peto's paradox revisited (revisited, revisited, revisited, and revisited yet again). *Proc. Natl. Acad. Sci. U. S. A.* **122**, e2502696122 (2025).
4. Tollis, M., Schneider-Utaka, A. K. & Maley, C. C. The evolution of human cancer gene duplications across mammals. *Mol. Biol. Evol.* **37**, 2875–2886 (2020).
